# Succinate mediates inflammation-induced adrenocortical dysfunction

**DOI:** 10.1101/2022.04.29.490066

**Authors:** Ivona Mateska, Anke Witt, Eman Hagag, Anupam Sinha, Canelif Yilmaz, Evangelia Thanou, Na Sun, Ourania Kolliniati, Maria Patschin, Heba Abdelmegeed, Holger Henneicke, Waldemar Kanczkowski, Ben Wielockx, Christos Tsatsanis, Andreas Dahl, Axel Walch, Ka Wan Li, Mirko Peitzsch, Triantafyllos Chavakis, Vasileia Ismini Alexaki

**Affiliations:** Institute of Clinical Chemistry and Laboratory Medicine, University Hospital, Technische Universität Dresden, Germany; Center of Neurogenomics and Cognitive Research (CNCR), Department of Molecular and Cellular Neurobiology, Vrije Universiteit, Amsterdam, Netherlands; Research Unit Analytical Pathology, German Research Center for Environmental Health, Helmholtz Zentrum München, Munich, Germany; Department of Clinical Chemistry, Medical School, University of Crete, Heraklion, Greece; Center for Regenerative Therapies, TU Dresden, Technische Universität Dresden, Germany; Department of Medicine III & Center for Healthy Ageing, Technische Universität Dresden, Germany; DRESDEN-concept Genome Center, Center for Molecular and Cellular Bioengineering, Technische Universität Dresden, Germany

**Author notes:** **Correspondence**: Dr. V.I. Alexaki, Institute for Clinical Chemistry and Laboratory Medicine, University Hospital, Technische Universität Dresden, Fetscherstrasse 74, 01307 Dresden, Germany, Dr. I. Mateska, Institute for Clinical Chemistry and Laboratory Medicine, University Hospital, Technische Universität Dresden, Fetscherstrasse 74, 01307 Dresden, Germany.

**Keywords:** adrenal gland, LPS, IL-1β, DNMT1, Succinate dehydrogenase, succinate, glucocorticoids

## Abstract

The hypothalamus-pituitary-adrenal (HPA) axis is activated in response to inflammation leading to increased production of anti-inflammatory glucocorticoids by the adrenal cortex, thereby representing an endogenous feedback loop. However, severe inflammation reduces the responsiveness of the adrenal gland to adrenocorticotropic hormone (ACTH) although the underlying mechanisms are poorly understood. Here, we show by transcriptomic, proteomic and metabolomic analyses that LPS-induced systemic inflammation triggers profound metabolic changes in steroidogenic adrenocortical cells, including downregulation of the TCA cycle and oxidative phosphorylation. Inflammation disrupts the TCA cycle at the level of succinate dehydrogenase (SDH) leading to succinate accumulation and disturbed steroidogenesis. Mechanistically, IL-1β reduces SDHB expression through upregulation of DNA methyltransferase 1 (DNMT1) and methylation of the SDHB promoter. Consequently, increased succinate levels impair oxidative phosphorylation and ATP synthesis, leading to reduced steroidogenesis. Together, we demonstrate that the IL-1β-DNMT1-SDHB-succinate axis disrupts steroidogenesis. Our findings not only provide a mechanistic explanation for the adrenal dysfunction in severe inflammation but also a potential target for therapeutic intervention.

## Introduction

Stress triggers the hypothalamic–pituitary–adrenal (HPA) axis, i.e. the release of corticotropin-releasing hormone (CRH) from the hypothalamus, followed by adrenocorticotropic hormone (ACTH) secretion from the anterior pituitary, which stimulates the synthesis of glucocorticoid hormones in the adrenal cortex, primarily cortisol in humans and corticosterone in rodents [1–3]. Similarly to any other stress stimulus, inflammation activates the HPA axis leading to increased glucocorticoid release, which is required to restrain the inflammatory response [4–8]. Adrenalectomized rodents show increased mortality after induction of systemic inflammation, while glucocorticoid administration increases survival [9,10]. Essentially, severe inflammation in sepsis is associated with impaired adrenal gland function [11–15] but the mechanisms remain poorly understood.

In immune cells, such as macrophages, dendritic cells and T cells, inflammation triggers cellular metabolic reprograming, enabling the cells to meet the increased demands for fast energy supply and anabolic processes [16–18]. How inflammation may affect cellular metabolism in other cell types and how this affects their function is less explored. Here we show that LPS-induced inflammation profoundly changes the cellular metabolism of steroidogenic adrenocortical cells, perturbing the TCA cycle at the level of succinate dehydrogenase B (SDHB). This is coupled to succinate accumulation, which impairs oxidative phosphorylation and leads to reduced steroidogenesis. Mechanistically, IL-1β inhibits *SDHB* expression through DNA methyltransferase 1 (DNMT1)-dependent DNA methylation of the *SDHB* promoter.

## Methods

### Animal experiments

Eight to twelve-week-old male C57BL/6J were injected i.p. with 1 mg/kg LPS (LPS-EB Ultrapure; InVivoGen) or PBS, and sacrificed after 6 h (for gene expression analyses) or 24 h (for all other analyses). *Irg1*^−/−^ and littermate control mice were injected with 3 mg/kg LPS and sacrificed after 16 h.

### Laser capture microdissection of adrenal cortex

Adrenal glands frozen in liquid nitrogen were cut in 25 – 30 μm thick sections, mounted on polyethylene naphthalate (PEN) membrane slides (Zeiss), dehydrated in increasing concentrations of ice-cold ethanol (75 %, 95 %, 100 %) for 45 sec each, and air-dried at room temperature (RT). Laser Capture Microdissection was performed with a Zeiss PALM MicroBeam LCM system. The adrenal cortex from 8 to 12 sections was microdissected and the tissue was collected on Adhesive Caps (Zeiss).

### Bioinformatic analysis of RNA-Seq data

For transcriptome mapping, strand-specific paired-end sequencing libraries from total RNA were constructed using TruSeq stranded Total RNA kit (Illumina Inc). Sequencing was performed on an Illumina HiSeq3000 (1×75 basepairs). Low quality nucleotides were removed with the Illumina fastq filter (http://cancan.cshl.edu/labmembers/gordon/fastq_illumina_filter/) and reads were further subjected to adaptor trimming using cutadapt [19]. Alignment of the reads to the Mouse genome was done using STAR Aligner [20] using the parameters: “–runMode alignReads –outSAMstrandField intronMotif –outSAMtype BAM SortedByCoordinate --readFilesCommand zcat”. Mouse Genome version GRCm38 (release M12 GENCODE) was used for the alignment. The parameters: ‘htseq-count -f bam -s reverse -m union -a 20’, HTSeq-0.6.1p1 [21] were used to count the reads that map to the genes in the aligned sample files. The GTF file (gencode.vM12.annotation.gtf) used for read quantification was downloaded from Gencode (https://www.gencodegenes.org/mouse/release_M12.html). Gene centric differential expression analysis was performed using DESeq2_1.8.1 [22]. The raw read counts for the genes across the samples were normalized using ‘rlog’ command of DESeq2 and subsequently these values were used to render a PCA plot using ggplot2_1.0.1 [23].

Pathway and functional analyses were performed using GSEA [24] and EGSEA [25]. GSEA is a stand-alone software with a GUI. To run GSEA, a ranked list of all the genes from DESeq2 based calculations was created by taking the −log10 of the p-value and multiplying it with the sign the of the fold change. This ranked list was then queried against Molecular Signatures Database (MsigDB), Reactome, KEGG and GO based repositories. EGSEA is an R/Bioconductor based command-line package. For doing functional analyses using EGSEA a differentially expressed list of genes with parameters log2foldchange > 0.3 and padj < 0.05 was used. Same database repositories as above were used for performing the functional analyses. For constructing pathway-specific heatmaps, the “rlog-normalized” expression values of the significantly expressed genes (padj < 0.05) were mapped on to the KEGG and GO pathways. These pathway-specific expression matrices were then scaled using z-transformation. The resulting matrices were visually rendered using MORPHEUS.

### Cell sorting

The adrenal cortex was separated from the medulla under a dissecting microscope and was digested in 1.6 mg/ml collagenase I (Sigma-Aldrich) and 1.6 mg/ml BSA in PBS, for 25 min at 37 °C while shaking at 900 rpm. The dissociated tissue was passed through a 22 G needle and 100 μm cell strainer and centrifuged at 300 g for 5 min at 4 °C. The cell suspension was washed in MACS buffer (0.5 % BSA, 2 mM EDTA in PBS) and CD31^+^ and CD45^+^ cells were sequentially positively selected using anti-CD31 and anti-CD45 MicroBeads (Miltenyi Biotec), respectively, according to manufacturer’s instructions. Briefly, pelleted cells resuspended in 190 μl MACS buffer, were mixed with 10 μl anti-CD31 MicroBeads, incubated for 15 min at 4 °C, washed with 2 ml MACS buffer and centrifuged at 300 g for 10 min at 4 °C. Then, the cell pellet was resuspended in 500 μl MACS buffer, applied onto MS Column placed on MACS Separator, and the flow-through (CD31^−^ cells) was collected. CD31^+^ cells were positively sorted from the MS Columns. The flow-through was centrifuged at 300 g for 5 min at 4 °C, and the pelleted cells were subjected to the same procedure using anti-CD45 MicroBeads, collecting the flow-through containing CD31^−^CD45^−^ adrenocortical cells. CD45^+^ cells were positively sorted from the MS Columns.

### MS/MS Proteomic analysis

CD31^−^CD45^−^ adrenocortical cells were sorted and snap-frozen. Samples were randomized and a gel-based sample preparation protocol was followed [26]. In brief, cell pellets were resuspended in SDS loading buffer and 30% acrylamide, boiled at 98 °C for 6 min, and 5 μg protein per sample were separated in 10% SDS gels (SurePAGE Bis-Tris gels, GenScript) for approximately 10 min at 120 V. The gels were fixed in 50 % (v/v) ethanol and 3 % (v/v) phosphoric acid and briefly stained with Colloidal Coomassie Blue. Sample containing lanes were sliced and cut into blocks of approximately 1 mm^3^, destained in 50 mM NH_4_HCO_3_ and 50 % (v/v) acetonitrile, dehydrated using 100 % acetonitrile, and rehydrated in 50 mM NH_4_HCO_3_ containing 10 μg / ml trypsin (sequence grade; Promega). After incubation overnight at 37 °C peptides were extracted and collected in a new tube, dried using a SpeedVac (Eppendorf), and stored at −20 °C until LC-MS analysis. Peptides were dissolved in 0.1 % formic acid, and 75 ng were loaded into EvoTips (EV2003, EvoSep) and washed according to manufacturer’s guidelines. The samples were run on a 15 cm x 75 μm, 1.9 μm Performance Column (EV1112, EvoSep) using the Evosep one liquid chromatography system with the 30 samples per day program. Peptides were analyzed by the TimsTof pro2 mass spectrometer (Bruker) with the diaPASEF method [27]. Data were analyzed using DIA-NN. The fasta database used was uniport mouse_UP000000589_10090. Deep learning was used to generate the in silico spectral library. Output was filtered at 0.01 FDR [28]. The Mass Spectrometry Downstream Analysis Pipeline (MS-DAP) (version beta 0.2.5.1) (https://github.com/ftwkoopmans/msdap)was used for quality control and candidate discovery [29]. Differential abundance analysis between groups was performed on log transformed protein abundances. Empirical Bayes moderated t-statistics with multiple testing correction by FDR, as implemented by the eBayes functions from the limma R package, was used as was previously described [30].

### Bioinformatics analysis of proteomics data

From the proteomics data, the missing data was imputed using the “impute” command running of “DEP” [31] package in R/Bioconductor [32] environment. The imputation was performed using “knn” function. The resultant imputed matrix was used for further analyses. Pathway and functional analyses were performed using GSEA [23] and EGSEA [24]. CLI version of GSEA v4.1 was run using the imputed matrix. Different pathway sets from Molecular Signatures Database (MsigDB) v7.2 like, HALLMARK, Biocarta, Reactome, KEGG, GO and WIKIPATHWAYS were queried for functional enrichment. Geneset permutations were performed 1,000 times to calculate the different statistical parameters. For doing functional analyses using EGSEA, imputed matrix was used. Same database repositories as above were used for performing the functional analyses.

### Quantitative RT – PCR

Total RNA was isolated from frozen adrenal glands with the TRI Reagent (MRC) after mechanical tissue disruption, extracted with chloroform and the NucleoSpin RNA Mini kit (Macherey-Nagel). Total RNA from sorted cells was isolated with the Rneasy Plus Micro Kit (Qiagen) according to manufacturer’s instructions. cDNA was synthesized with the iScript cDNA Synthesis kit (Biorad) and gene expression was determined using the SsoFast Eva Green Supermix (Bio-Rad), with a CFX384 real-time System C1000 Thermal Cycler (Bio-Rad) and the Bio-Rad CFX Manager 3.1 software. The relative gene expression was calculated using the ΔΔCt method, *18S* was used as a reference gene. Primers are listed in Table 5.

**Table 1.**
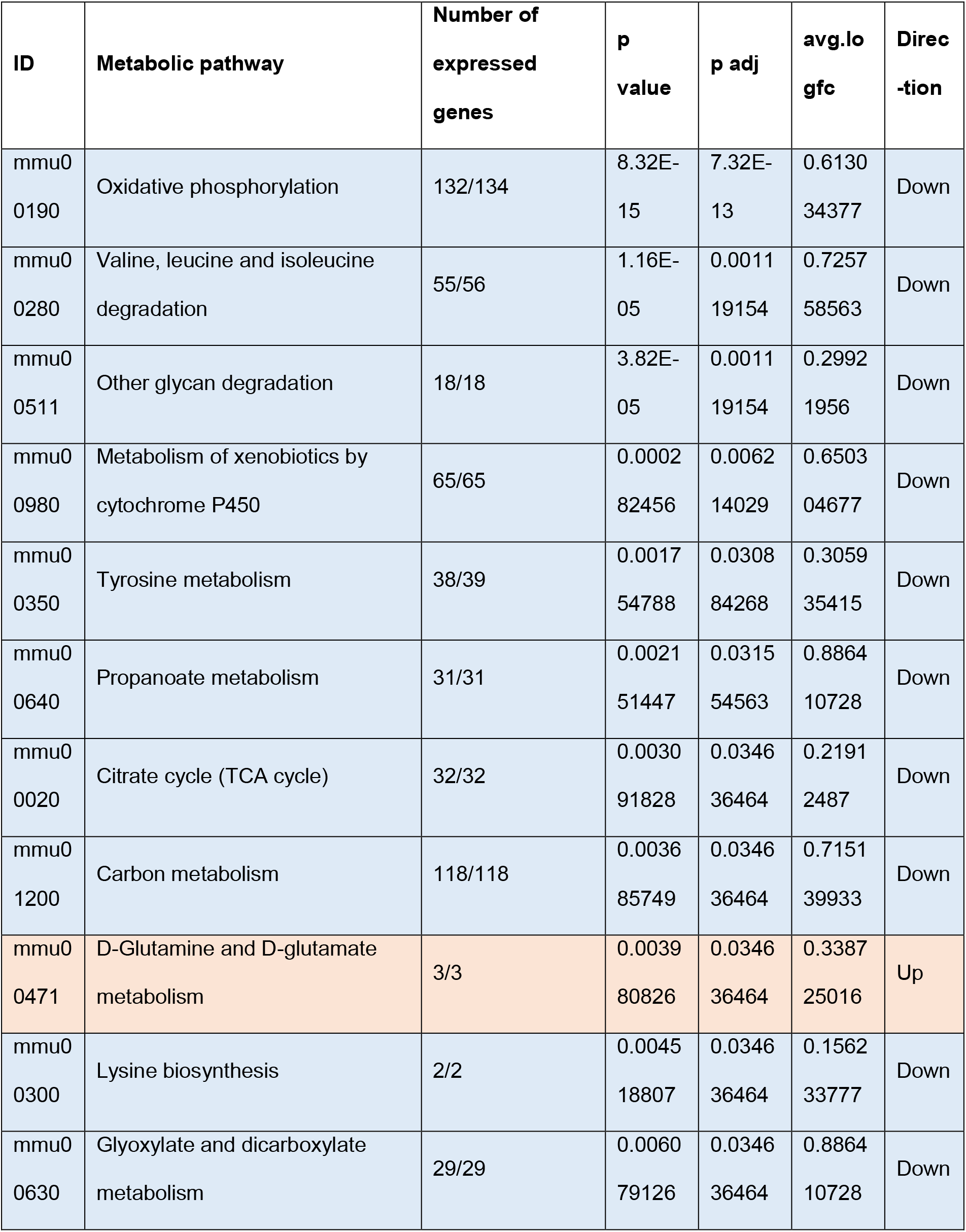
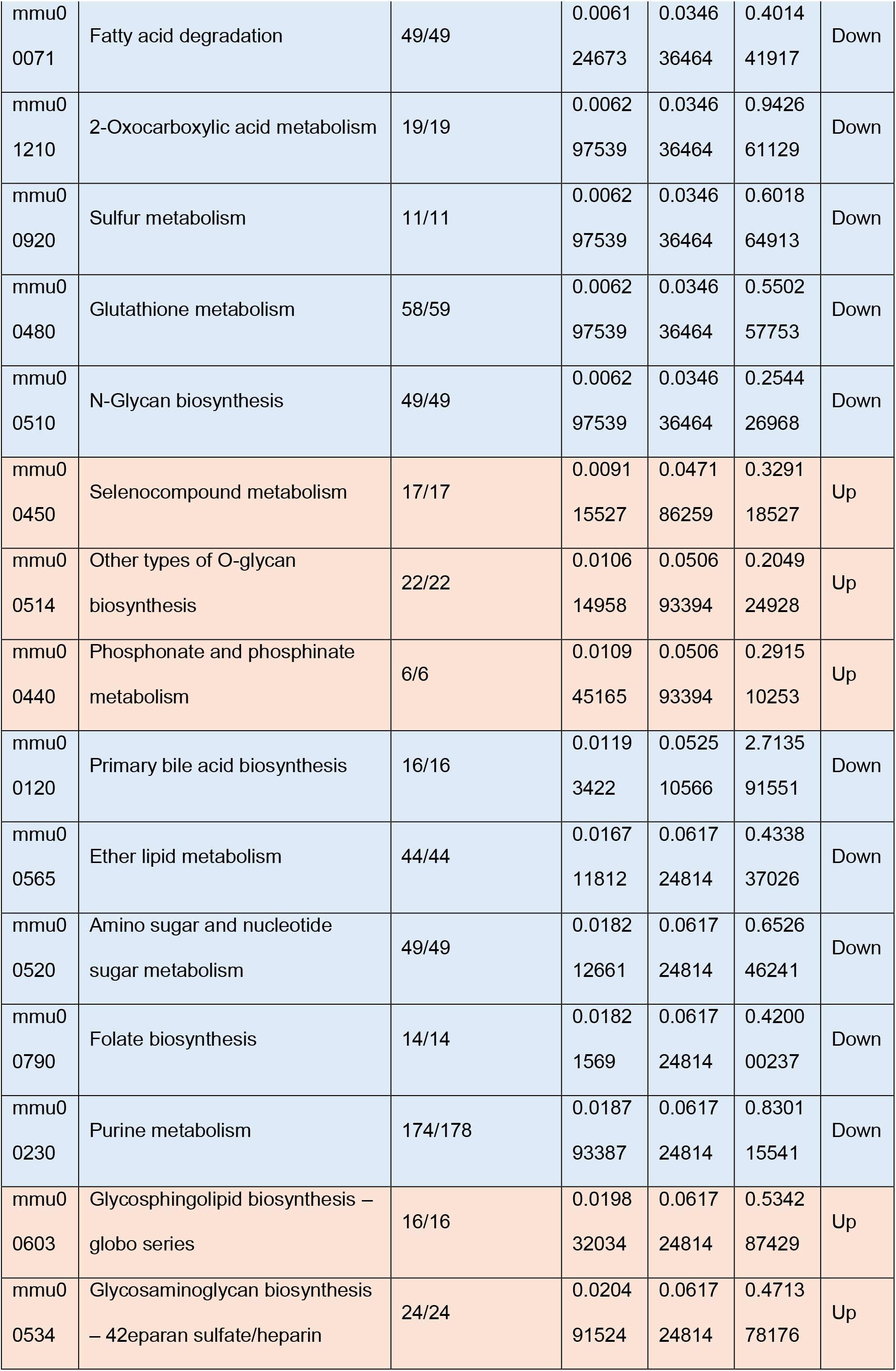
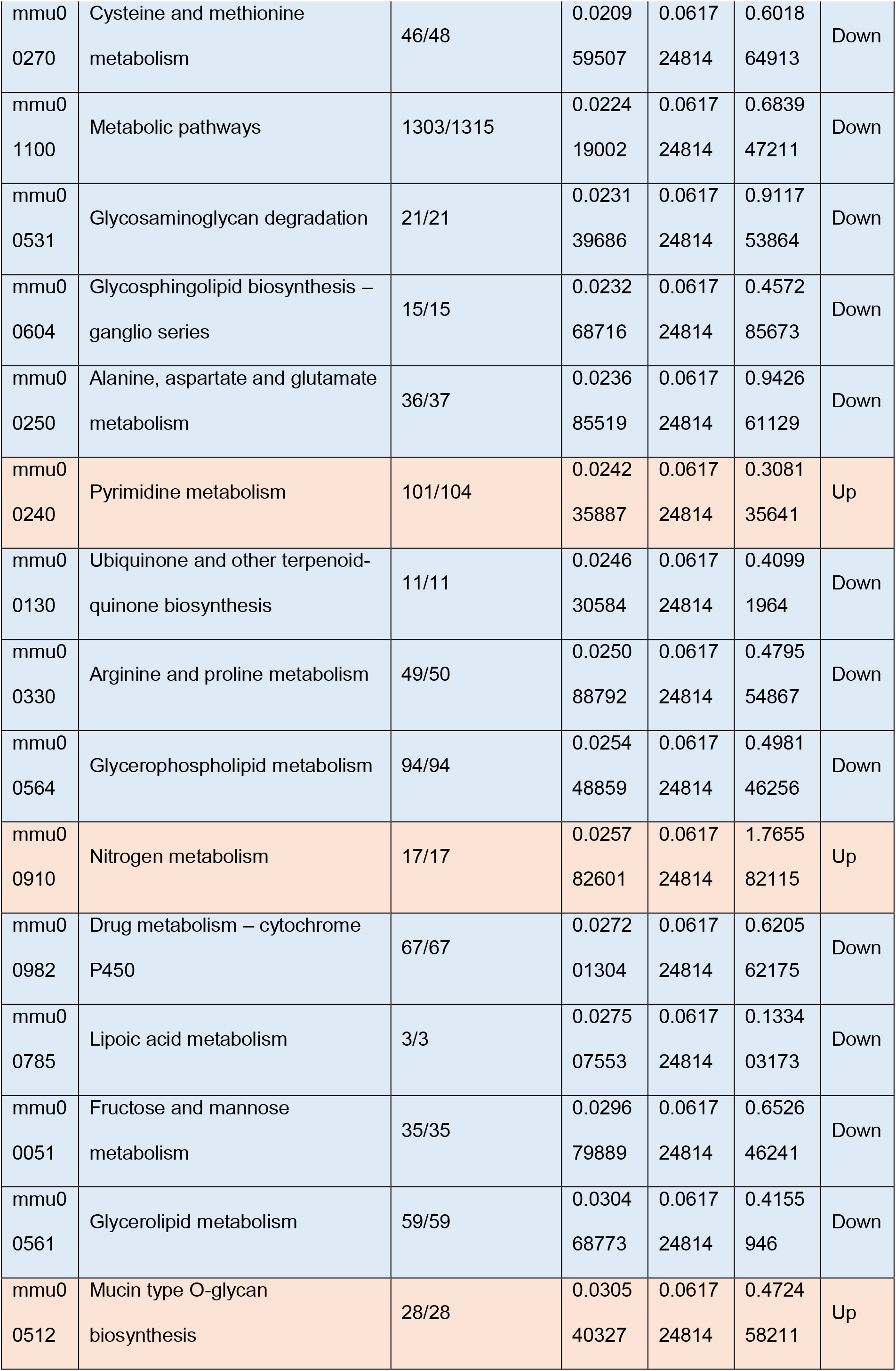
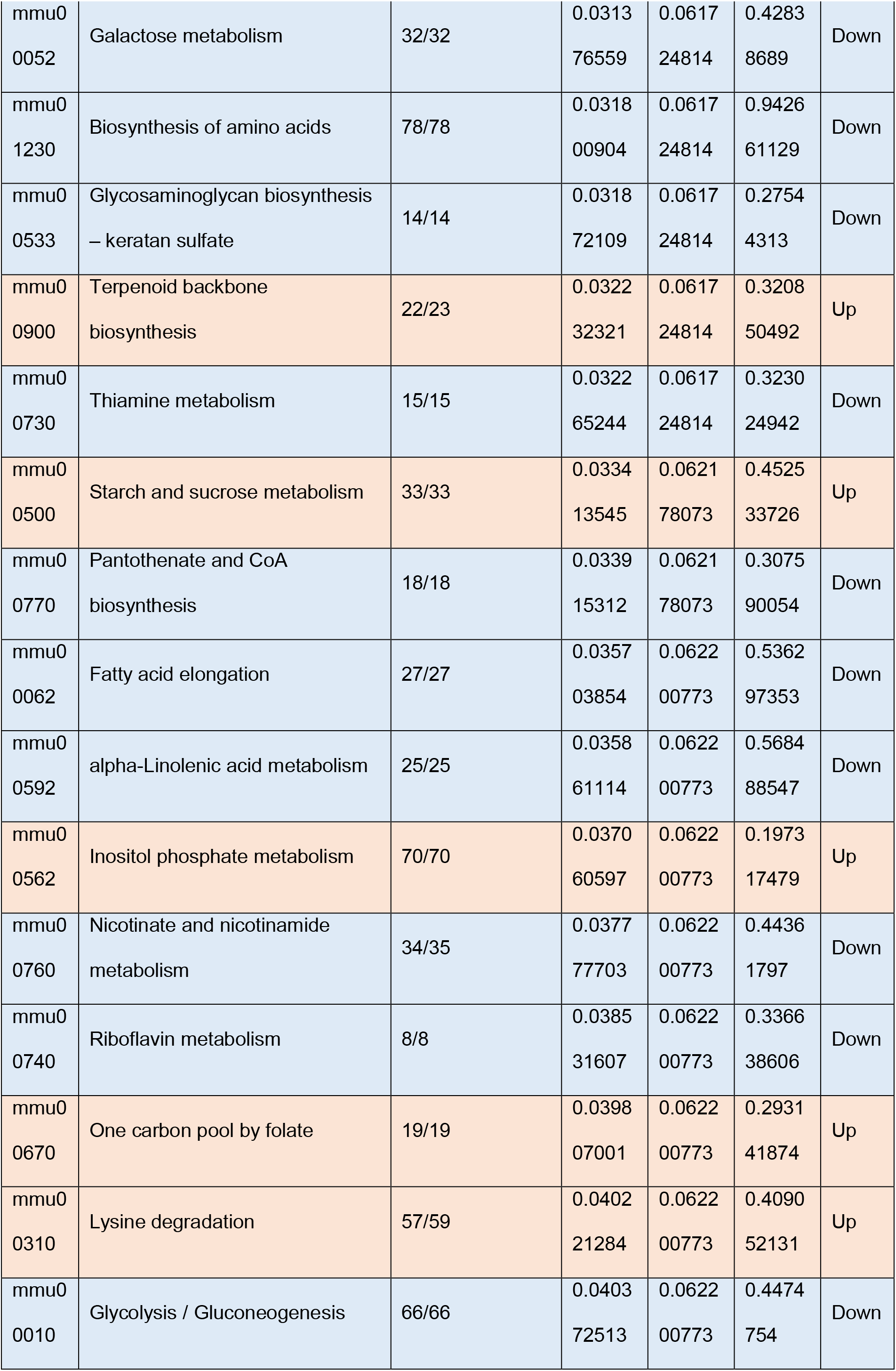
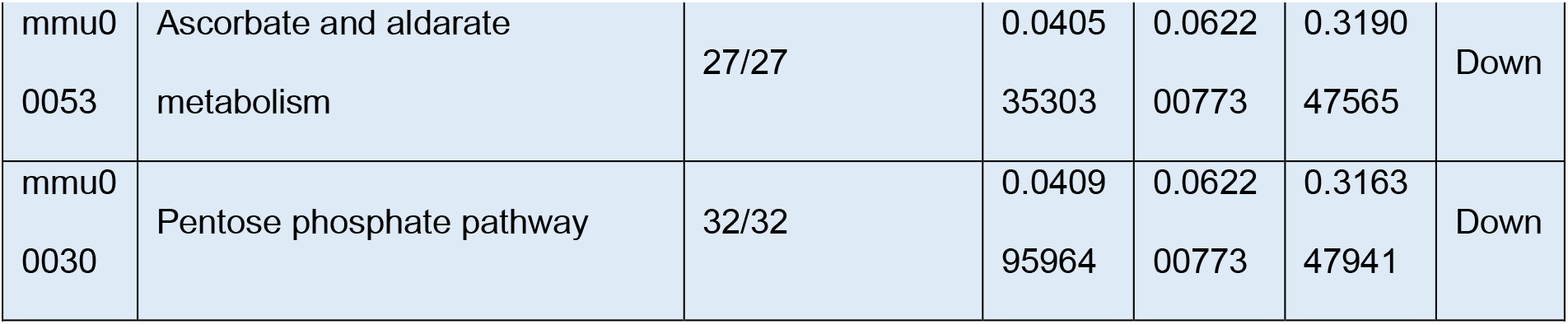
Cellular metabolic pathways transcriptionally regulated by inflammation in the adrenal cortex. The pathway analysis of differentially expressed genes was done with the software package EGSEA and queried against the KEGG Pathways repository. Pathways with p < 0.05 are shown.

**Table 2.** Cellular metabolic pathways regulated on protein level by inflammation in adrenocortical cells. The pathway analysis of differentially expressed proteins was done with the software package EGSEA and queried against the KEGG Pathways repository. Pathways with p < 0.05 are shown.

| ID | Metabolic pathway | Number of detected proteins | p.value | p.adj | avg.logfc | Direction |
| --- | --- | --- | --- | --- | --- | --- |
| mmu00534 | Glycosaminoglycan biosynthesis – heparan sulfate / heparin | 4/24 | 1.50E-07 | 1.29E-05 | 0.13 | Up |
| mmu00280 | Valine, leucine and isoleucine degradation | 37/57 | 5.20E-07 | 2.23E-05 | 0.01 | Down |
| mmu00983 | Drug metabolism – other enzymes | 25/92 | 3.80E-05 | 0.000856142 | 0.02 | Down |
| mmu00562 | Inositol phosphate metabolism | 19/72 | 5.89E-05 | 0.000856142 | 0.03 | Up |
| mmu00260 | Glycine, serine and threonine metabolism | 19/40 | 6.89E-05 | 0.000856142 | 0.03 | Down |
| mmu00240 | Pyrimidine metabolism | 25/58 | 6.98E-05 | 0.000856142 | 0.04 | Up |
| mmu00982 | Drug metabolism – cytochrome P450 | 16/71 | 7.76E-05 | 0.000856142 | 0.02 | Down |
| mmu00230 | Purine metabolism | 44/133 | 7.96E-05 | 0.000856142 | 0.04 | Up |
| mmu0<br>0790 | Folate biosynthesis | 9/26 | 9.75E-05 | 0.0009<br>31452 | 0.09 | Up |
| mmu0<br>1100 | Metabolic pathways | 593/1608 | 0.0001<br>34694 | 0.0011<br>51176 | 0.03 | Down |
| mmu0<br>0190 | Oxidative phosphorylation | 70/135 | 0.0001<br>60368 | 0.0011<br>51176 | 0.02 | Down |
| mmu0<br>0980 | Metabolism of xenobiotics by<br>cytochrome P450 | 19/73 | 0.0001<br>75997 | 0.0011<br>51176 | 0.02 | Down |
| mmu0<br>1240 | Biosynthesis of cofactors | 74/154 | 0.0001<br>78113 | 0.0011<br>51176 | 0.04 | Down |
| mmu0<br>0730 | Thiamine metabolism | 5/15 | 0.0001<br>87401 | 0.0011<br>51176 | 0.02 | Down |
| mmu0<br>0020 | Citrate cycle (TCA cycle) | 30/32 | 0.0002<br>0177 | 0.0011<br>56812 | 0.01 | Down |
| mmu0<br>1200 | Carbon metabolism | 81/121 | 0.0004<br>04805 | 0.0021<br>75826 | 0.02 | Down |
| mmu0<br>0760 | Nicotinate and nicotinamide<br>metabolism | 11/41 | 0.0004<br>59649 | 0.0023<br>25283 | 0.04 | Up |
| mmu0<br>0860 | Porphyrin and chlorophyll<br>metabolism | 19/43 | 0.0005<br>49279 | 0.0026<br>24333 | 0.04 | Up |
| mmu0<br>0360 | Phenylalanine metabolism | 7/23 | 0.0007<br>44591 | 0.0033<br>70253 | 0.01 | Up |
| mmu0<br>0630 | Glyoxylate and dicarboxylate<br>metabolism | 20/32 | 0.0007<br>94408 | 0.0034<br>15956 | 0.02 | Down |
| mmu0<br>0480 | Glutathione metabolism | 28/72 | 0.0010<br>37994 | 0.0042<br>50832 | 0.02 | Down |
| mmu0<br>0061 | Fatty acid biosynthesis | 12/19 | 0.0011<br>9934 | 0.0043<br>23466 | 0.04 | Up |
| mmu0<br>0052 | Galactose metabolism | 17/32 | 0.0012<br>12032 | 0.0043<br>23466 | 0.02 | Down |
| mmu0<br>0350 | Tyrosine metabolism | 12/40 | 0.0012<br>75764 | 0.0043<br>23466 | 0.02 | Down |
| mmu0<br>0900 | Terpenoid backbone biosynthesis | 8/23 | 0.0012<br>83046 | 0.0043<br>23466 | 0.01 | Down |
| mmu0<br>0140 | Steroid hormone biosynthesis | 12/92 | 0.0013<br>07094 | 0.0043<br>23466 | 0.02 | Down |
| mmu0<br>0511 | Other glycan degradation | 11/18 | 0.0016<br>20057 | 0.0051<br>60182 | 0.03 | Down |
| mmu0<br>0520 | Amino sugar and nucleotide sugar metabolism | 29/51 | 0.0019<br>22509 | 0.0059<br>04848 | 0.02 | Down |
| mmu0<br>0040 | Pentose and glucuronate interconversions | 9/35 | 0.0022<br>91756 | 0.0067<br>96243 | 0.02 | Down |
| mmu0<br>0620 | Pyruvate metabolism | 34/44 | 0.0047<br>41112 | 0.0134<br>28366 | 0.02 | Down |
| mmu0<br>0524 | Neomycin, kanamycin and gentamicin biosynthesis | 3/5 | 0.0048<br>40458 | 0.0134<br>28366 | 0.02 | Down |
| mmu0<br>0053 | Ascorbate and aldarate metabolism | 9/31 | 0.0052<br>20665 | 0.0134<br>29037 | 0.01 | Down |
| mmu0<br>0830 | Retinol metabolism | 8/97 | 0.0052<br>79453 | 0.0134<br>29037 | 0.01 | Down |
| mmu0<br>0531 | Glycosaminoglycan degradation | 10/21 | 0.0053<br>09154 | 0.0134<br>29037 | 0.02 | Down |
| mmu0<br>0450 | Selenocompound metabolism | 9/17 | 0.0069<br>51355 | 0.0170<br>80471 | 0.02 | Down |
| mmu0<br>0250 | Alanine, aspartate and glutamate metabolism | 17/39 | 0.0079<br>55263 | 0.0190<br>04239 | 0.02 | Down |
| mmu0<br>1230 | Biosynthesis of amino acids | 45/79 | 0.0088<br>73164 | 0.0206<br>24111 | 0.02 | Down |
| mmu0<br>1212 | Fatty acid metabolism | 40/62 | 0.0093<br>36325 | 0.0211<br>29578 | 0.03 | Down |
| mmu0<br>0500 | Starch and sucrose metabolism | 14/34 | 0.0111<br>70016 | 0.0246<br>31316 | 0.02 | Down |
| mmu0<br>0514 | Other types of O-glycan biosynthesis | 15/43 | 0.0118<br>76496 | 0.0253<br>49557 | 0.04 | Up |
| mmu0<br>1210 | 2-Oxocarboxylic acid metabolism | 11/20 | 0.0122<br>7639 | 0.0253<br>49557 | 0.01 | Down |
| mmu0<br>0650 | Butanoate metabolism | 13/28 | 0.0124<br>04185 | 0.0253<br>49557 | 0.01 | Down |
| mmu0<br>0670 | One carbon pool by folate | 9/19 | 0.0126<br>74778 | 0.0253<br>49557 | 0.05 | Down |
| mmu0<br>0310 | Lysine degradation | 20/64 | 0.0140<br>1052 | 0.0273<br>84197 | 0.02 | Down |
| mmu0<br>0590 | Arachidonic acid metabolism | 9/86 | 0.0152<br>53177 | 0.0291<br>50517 | 0.01 | Down |
| mmu0<br>0770 | Pantothenate and CoA biosynthesis | 8/21 | 0.0166<br>32522 | 0.0310<br>95584 | 0.02 | Down |
| mmu0<br>0592 | alpha-Linolenic acid metabolism | 3/25 | 0.0178<br>19371 | 0.0326<br>05657 | 0.03 | Up |
| mmu0<br>0780 | Biotin metabolism | 3/3 | 0.0186<br>50597 | 0.0334<br>15653 | 0.01 | Down |
| mmu0<br>0640 | Propanoate metabolism | 25/31 | 0.0201<br>82799 | 0.0351<br>73564 | 0.02 | Down |
| mmu0<br>0290 | Valine, leucine and isoleucine biosynthesis | 2/4 | 0.0205<br>3428 | 0.0351<br>73564 | 0.02 | Down |
| mmu0<br>0920 | Sulfur metabolism | 7/11 | 0.0212<br>06387 | 0.0351<br>73564 | 0.02 | Down |
| mmu0<br>0062 | Fatty acid elongation | 15/19 | 0.0212<br>67736 | 0.0351<br>73564 | 0.02 | Down |
| mmu0<br>0604 | Glycosphingolipid biosynthesis – ganglio series | 5/15 | 0.0220<br>65753 | 0.0358<br>04807 | 0.02 | Down |
| mmu00563 | Glycosylphosphatidylinositol (GPI)-anchor biosynthesis | 8/26 | 0.0235<br>53511 | 0.0361<br>86307 | 0.04 | Down |
| mmu00750 | Vitamin B6 metabolism | 3/9 | 0.0235<br>53511 | 0.0361<br>86307 | 0.03 | Up |
| mmu00220 | Arginine biosynthesis | 7/20 | 0.0242<br>45413 | 0.0361<br>86307 | 0.02 | Down |
| mmu00270 | Cysteine and methionine metabolism | 29/53 | 0.0254<br>10276 | 0.0361<br>86307 | 0.03 | Up |
| mmu00051 | Fructose and mannose metabolism | 17/36 | 0.0255<br>1991 | 0.0361<br>86307 | 0.04 | Down |
| mmu00071 | Fatty acid degradation | 30/52 | 0.0256<br>67032 | 0.0361<br>86307 | 0.02 | Down |
| mmu00330 | Arginine and proline metabolism | 23/54 | 0.0256<br>67032 | 0.0361<br>86307 | 0.02 | Down |
| mmu00561 | Glycerolipid metabolism | 23/62 | 0.0256<br>67032 | 0.0361<br>86307 | 0.03 | Down |
| mmu00010 | Glycolysis / Gluconeogenesis | 40/67 | 0.0507<br>28869 | 0.0681<br>66917 | 0.02 | Down |
| mmu00030 | Pentose phosphate pathway | 17/33 | 0.0507<br>28869 | 0.0681<br>66917 | 0.03 | Down |
| mmu00410 | beta-Alanine metabolism | 19/32 | 0.0507<br>28869 | 0.0681<br>66917 | 0.01 | Down |

**Table 3.** ROS pathways are transcriptionally upregulated in the adrenal cortex of LPS-treated mice. The pathway analysis of differentially expressed genes was done with the software package EGSEA and queried against the GO Gene Sets repository. Pathways with padj. < 0.05 are shown.

| ID | Gene Set | Number of<br>expressed<br>genes | p<br>value | p adj | avg.l<br>ogfc | Direc<br>-tion |
| --- | --- | --- | --- | --- | --- | --- |
| M13<br>446 | GO Regulation of reactive oxygen<br>species metabolic process | 271/275 | 3.75E-08 | 1.26E-06 | 1.01<br>00 | Up |
| M13<br>580 | GO Positive regulation of reactive<br>oxygen species metabolic process | 182/186 | 2.33E-07 | 6.26E-06 | 1.01<br>00 | Up |
| M16<br>953 | GO Response to reactive oxygen<br>species | 300/317 | 0.0034<br>22132 | 0.0090<br>35375 | 0.86<br>00 | Up |
| M16<br>581 | GO Cellular response to reactive<br>oxygen species | 173/177 | 0.0025<br>37297 | 0.0090<br>35375 | 0.76<br>00 | Up |
| M10<br>618 | GO Negative regulation of response to<br>reactive oxygen species | 24/24 | 0.0072<br>384 | 0.0109<br>42115 | 0.71<br>00 | Up |
| M15<br>379 | GO Regulation of reactive oxygen<br>species biosynthetic process | 145/148 | 5.90E-05 | 0.0007<br>70609 | 0.70<br>00 | Up |
| M10<br>827 | GO Positive regulation of reactive<br>oxygen species biosynthetic process | 120/123 | 0.0004<br>65606 | 0.0045<br>4261 | 0.70<br>00 | Up |
| M15<br>990 | GO Reactive oxygen species<br>metabolic process | 163/167 | 0.0089<br>36465 | 0.0125<br>68942 | 0.67<br>00 | Up |
| M16<br>764 | GO Regulation of response to reactive<br>oxygen species | 43/43 | 0.0067<br>53207 | 0.0104<br>98628 | 0.62<br>00 | Down |
| M16<br>007 | GO Negative regulation of reactive<br>oxygen species biosynthetic process | 23/23 | 0.0164<br>34387 | 0.0202<br>74996 | 0.60<br>00 | Up |
| M12<br>185 | GO Reactive oxygen species<br>biosynthetic process | 32/33 | 0.0064<br>83259 | 0.0102<br>4232 | 0.57<br>00 | Down |
| M10<br>894 | GO Negative regulation of reactive<br>oxygen species metabolic process | 59/59 | 0.0065<br>38654 | 0.0102<br>87661 | 0.55<br>00 | Up |

**Table 4.** ROS – related protein expression is upregulated in the adrenal cortex of LPS-treated mice. The pathway analysis of differentially expressed proteins was done with the software package EGSEA and queried against the GO Gene Sets repository. Pathways with padj. < 0.05 are shown.

| ID | Protein Set | Number of detected proteins | p.value | p.adj | avg.logfc | Direction |
| --- | --- | --- | --- | --- | --- | --- |
| M13<br>446 | GO REGULATION OF REACTIVE OXYGEN SPECIES METABOLIC PROCESS | 62/275 | 6.30E-06 | 0.000<br>12401 | 0.03 | Up |
| M15<br>379 | GO REGULATION OF REACTIVE OXYGEN SPECIES BIOSYNTHETIC PROCESS | 34/148 | 9.43E-06 | 0.000<br>14372 | 0.03 | Up |
| M13<br>580 | GO POSITIVE REGULATION OF REACTIVE OXYGEN SPECIES METABOLIC PROCESS | 29/186 | 1.11E-05 | 0.000<br>15778 | 0.03 | Up |
| M10<br>827 | GO POSITIVE REGULATION OF REACTIVE OXYGEN SPECIES BIOSYNTHETIC PROCESS | 20/123 | 1.17E-05 | 0.000<br>16203 | 0.03 | Up |
| M10<br>894 | GO NEGATIVE REGULATION OF REACTIVE OXYGEN SPECIES METABOLIC PROCESS | 23/59 | 5.29E-05 | 0.000<br>37856 | 0.05 | Down |
| M16<br>007 | GO NEGATIVE REGULATION OF REACTIVE OXYGEN SPECIES BIOSYNTHETIC PROCESS | 12/23 | 0.00014<br>126 | 0.000<br>73167 | 0.07 | Up |
| M16<br>581 | GO CELLULAR RESPONSE TO REACTIVE OXYGEN SPECIES | 50/177 | 0.00015<br>378 | 0.000<br>77551 | 0.03 | Down |
| M15<br>990 | GO REACTIVE OXYGEN SPECIES METABOLIC PROCESS | 35/167 | 0.00037<br>436 | 0.001<br>50479 | 0.02 | Down |
| M16<br>953 | GO RESPONSE TO REACTIVE<br>OXYGEN SPECIES | 83/317 | 0.00076<br>076 | 0.002<br>62274 | 0.03 | Up |
| M12<br>185 | GO REACTIVE OXYGEN SPECIES<br>BIOSYNTHETIC PROCESS | 9/33 | 0.00192<br>844 | 0.005<br>45214 | 0.02 | Down |

**Table 5.**
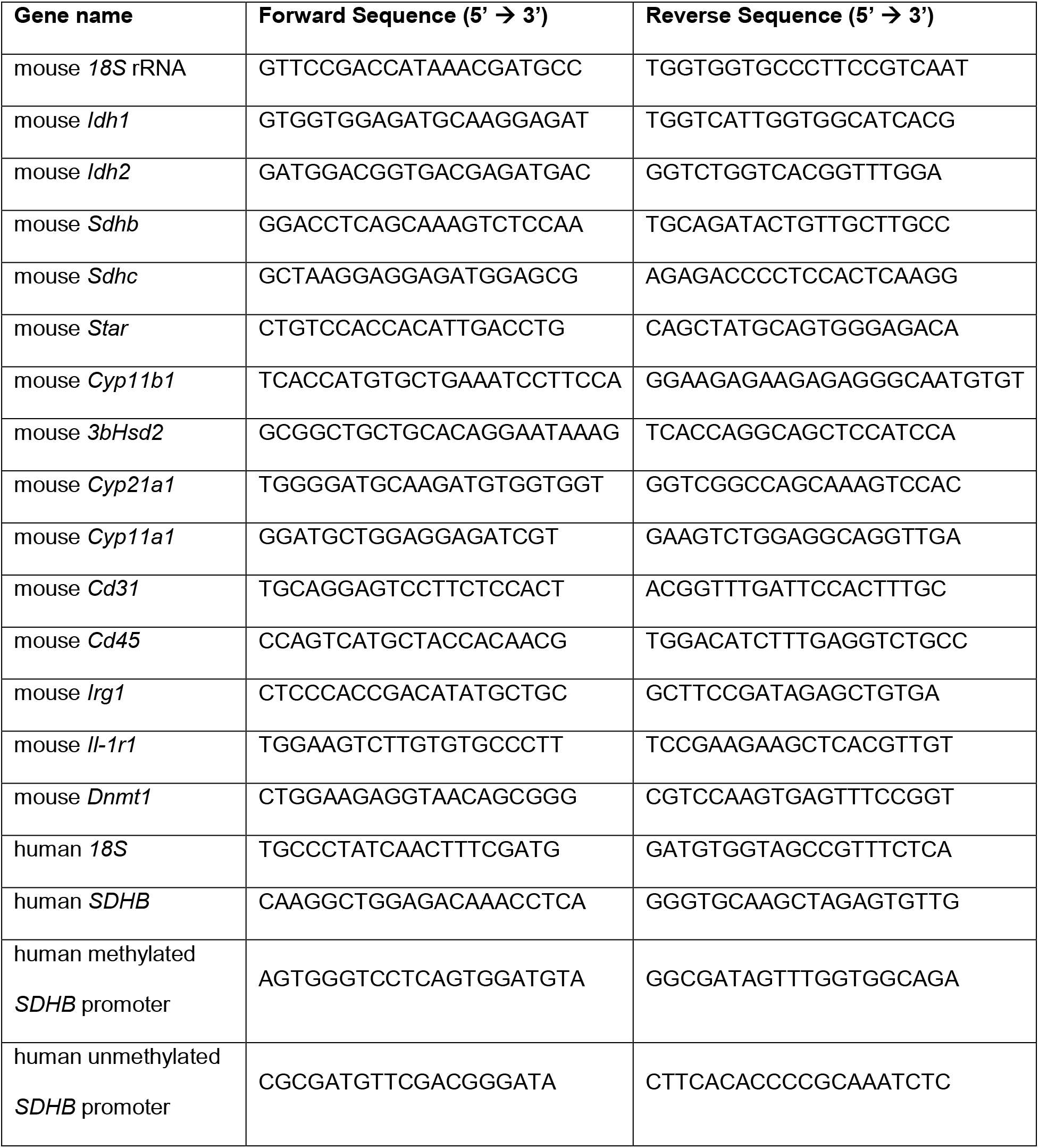
Primer sequences.

### Cell culture and in vitro treatments

CD31^−^CD45^−^ adrenocortical cells were plated on 0.2 % gelatin-coated wells of 96-well plates in DMEM/F12 medium supplemented with 1 % fetal bovine serum (FBS), 50 U/mL penicillin and 50 μg/mL streptomycin (all from Gibco), and let to attach for an hour before treatments. Cells from both adrenal glands from each mouse were pooled together and plated in 2 wells of a 96-well plate. Mouse adrenal explants were dissected from surrounding fat and left in DMEM/F12 medium with 1 % FBS, 50 U/mL penicillin and 50 μg/mL streptomycin for an hour before treatments. NCI-H295R cells (obtained from ATCC) were maintained in DMEM/F12 medium supplemented with 2.5 % Nu-Serum type I (Corning), 1 % Insulin Transferrin Selenium (ITS; Gibco), 50 U/mL penicillin and 50 μg/mL streptomycin.

Cells or explants were treated with DMM (20 mM; Sigma-Aldrich), DES (5 mM; Sigma-Aldrich), FCCP (1 μM; Agilent Technologies), OM (500 nM; Agilent Technologies), AG-221 (10 μM; Selleckchem), 4-OI (125 μM; Cayman Chemical), IL-1β(20 ng/ml, PeproTech), IL-6 (20 ng/ml, PeproTech), TNFα(20 ng/ml, PeproTech), LPS (1 μg/ml; InVivoGen), ACTH (100 ng/ml; Sigma-Aldrich) or Forskolin (10 μM; Sigma-Aldrich). siRNA transfections were done with ON-TARGETplus SMARTpool siRNA against *SDHB* (10 nM), *Sdhb* (30 nM), *Idh2* (30 nM) or *Dnmt1* (30 nM) (all from Horizon Discovery), with Lipofectamine RNAiMAX transfection reagent (Invitrogen), using a reverse transfection protocol per manufacturer’s instructions.

### Steroid hormone measurement

Steroid hormones were analyzed by LC-MS/MS in cell culture or explant supernatants as described previously [33]. Fifty to hundred μL cell culture supernatants were extracted by solid phase extraction using positive pressure, followed by a dry-down under gentle stream of nitrogen. Residues were reconstituted in 100 μL of the initial LC mobile phase and 10 μL were injected for detection by the triple quadrupole mass spectrometer in multiple reaction-monitoring scan mode using positive atmospheric pressure chemical ionization. Quantification of steroid concentrations was done by comparisons of ratios of analyte peak areas to respective peak areas of stable isotope labeled internal standards obtained in samples to those of calibrators.

### Measurement of TCA cycle metabolites

TCA cycle metabolites were determined by LC-MS/MS as described before [34]. Itaconate was included in the existing LC-MS/MS method using multi-reaction monitoring (MRM)-derived ion transition of 128.9->85.1. For quantification of itaconate ratios of analyte peak areas to respective peak areas of the stable isotope labeled internal standard (itaconic acid-^13^C5; Bio-Connect B.V., The Netherlands; MRM transition 133.9->89.1) obtained in samples were compared to those of calibrators.

### MALDI-FT-ICR-MSI

Tissue preparation steps for MALDI imaging mass spectrometry (MALDI-MSI) analysis was performed as previously described [35,36]. Frozen mouse adrenals were cryosectioned at 12 μm (CM1950, Leica Microsystems, Wetzlar, Germany) and thaw-mounted onto indium-tin-oxide coated conductive slides (Bruker Daltonik, Bremen, Germany). The matrix solution consisted of 10 mg/ml 1,5-Diaminonaphthalene (Sigma-Aldrich, Germany) in water/acetonitrile 30:70 (v/v). SunCollectTM automatic sprayer (Sunchrom, Friedrichsdorf, Germany) was used for matrix application. The MALDI-MSI measurement was performed on a Bruker Solarix 7T FT-ICR-MS (Bruker Daltonik, Bremen, Germany) in negative ion mode using 100 laser shots at a frequency of 1000 Hz. The MALDI-MSI data were acquired over a mass range of m/z 75-250 with 50 μm lateral resolution. Following the MALDI imaging experiments, the tissue sections were stained with hematoxylin and eosin (H&E) and scanned with an AxioScan.Z1 digital slide scanner (Zeiss, Jena, Germany) equipped with a 20x magnification objective. After the MALDI-MSI measurement, the acquired data underwent spectra processing in FlexImaging v. 5.0 (Bruker Daltonik, Bremen, Germany) and SciLS Lab v. 2021 (Bruker Daltonik, Bremen, Germany). The mass spectra were root-mean-square normalized. MS Peak intensity of isocitrate and succinate of adrenal cortex regions were exported and applied for relative quantification analysis.

### ATP measurement

Total ATP was measured in adrenal glands using the ATP Assay Kit (ab83355, Abcam). Briefly, adrenal glands were collected, washed with PBS and immediately homogenized in 100 μl ATP assay buffer. Samples were cleared using the Deproteinizing Sample Preparation Kit – TCA (ab204708, Abcam). Samples were incubated for 30 min with the ATP reaction mix and fluorescence (Ex / Em = 535 / 587 nm) was measured using the Synergy HT microplate reader. The recorded measurements were normalized to the weight of the adrenal gland.

### ROS measurement

ROS was detected using the DCFDA/H2DCFDA Cellular ROS Detection Assay Kit (ab113851, Abcam). NCI-H295R cells were plated at 80,000 cells / well in 96-well plate with black walls and clear bottom (Corning) and were incubated with 20 μM DCFDA Solution for 45 min at 37 °C in dark. Fluorescence (Ex / Em = 485 / 535 nm) was measured using the Synergy HT microplate reader.

### ADP / ATP ratio measurement

Intracellular ATP / ADP ratio was determined with the ADP / ATP Ratio Assay Kit (MAK135, Sigma-Aldrich). NCI-H295R cells were plated at 80,000 cells / well in 96-well plate with white flat bottom wells (Corning). Luminescence was measured using the Synergy HT microplate reader.

### NADP/NADPH measurement

Intracellular NADP/NADPH ratio was measured with the NADP/NADPH Assay Kit (Fluorometric) (ab176724, Abcam). NCI-H295R cells were plated at 5×10^6^ cells / 10 cm – diameter dish. Fluorescence (Ex / Em = 540 / 590 nm) was measured using the Synergy HT microplate reader. NADPH in adrenal tissue homogenates were analyzed by LC-MS/MS using an adapted method previously described [37].

### Enzyme activity measurement

SDH and IDH activities were measured using respective colorimetric assay kits (MAK197, Sigma-Aldrich, ab102528, Abcam). Cortices from both adrenal glands of each mouse were pooled and processed together. Absorbance (at 600 nm for SDH or 450 nm for IDH) was detected using the Synergy HT microplate reader.

### Seahorse assay

OCR measurements were performed with a Seahorse XF96 Analyzer (Agilent Technologies). NCI-H295R cells were plated at 80,000 cells / well in 0.2 % gelatin-precoated XF96 cell culture microplate (Agilent). The experimental medium used was XF Base Medium supplemented with glucose (10 mM), pyruvate (1 mM) and glutamine (2 mM).

### Measurement of mitochondrial load and membrane potential

The adrenal cortex was digested and dissociated cells were incubated with Mitotracker Green (0.25 μM; Invitrogen), TMRE (2.5 μM; Thermofisher), CD31-PeCy7 (1:100; eBioscience) and CD45-PeCy7 (1:100; eBioscience) for 30 min in FACS buffer (0.5 % BSA, 2 mM EDTA in PBS) at 37°C in dark. Live cells were selected by Hoechst staining. NCI-H295R cells were incubated with MitotrackerGreen (100 nM) and TMRE (100 nM) for 30 min at 37 °C in dark. FACS was performed using LSR Fortessa X20 flow cytometer and data were analyzed with the FlowJo software.

### Western blotting

Cells were lysed with 10 mM Tris-HCl, pH7.4 + 1 % SDS + 1 mM sodium vanadate, cell lysates were centrifuged at 16,000 g for 5 min at 4 °C, supernatants were collected and total protein concentration was measured using Pierce BCA Protein Assay Kit (Thermo Scientific). Gel electrophoresis was performed according to standard protocols [38]. Protein samples were prepared with 5x Reducing Laemmli buffer, denatured at 95 °C for 5 min and loaded on a 10 % acrylamide gel (Invitrogen) for sodium dodecyl sulfate polyacrylamide gel electrophoresis (SDS-PAGE). PageRuler Prestained Protein Ladder (Thermo Fisher Scientific) was used as a protein size ladder. The separated proteins were transferred on Amersham Protran nitrocellulose membrane (GE Healthcare Lifescience). After blocking with 5 % skimmed milk in TBS-T (0.1 % Tween-20 (Sigma-Aldrich) in 1x Tris-buffered saline) for 1 hour at RT, membranes were incubated overnight at 4 °C with anti-SDHB (1:1000; Sigma-Aldrich, #HPA002868), anti-DNMT1 (1:1000; Cell Signaling, #5032) or anti-Tubulin (1:3000; Sigma-Aldrich, T5186), diluted in 5 % BSA in TBS-T. After washing, membranes were incubated for 1 hour at RT with secondary antibodies: goat anti-rabbit IgG HRP-conjugated (1:3000; Jackson ImmunoResearch) or goat anti-mouse IgG HRP-conjugated (1:3000; Jackson ImmunoResearch), diluted in 5 % skimmed milk in TBS-T. The signal was detected using the Western Blot Ultra-Sensitive HRP Substrate (Takara) and imaged using the Fusion FX Imaging system (PeqLab Biotechnologie).

### DNA methylation measurement

Genomic DNA from 2×10^6^ NCI-H295R cells was isolated with the Quick-DNA Miniprep Kit (Zymo research). Bisulfite treatment was performed using the EZ DNA Methylathion™ Kit (Zymo Research), following the manufacturer’s protocol. For each sample 500 ng genomic DNA was used, bisulfite treated for 14h in the dark and, after a desulphonation and cleaning step, eluted in 10 μL nuclease-free water. The SDHB promoter region was amplified with primers for a methylated and a non-methylated sequence (listed in Table 5), using the Qiagen Multiplex PCR Kit. Equal amount of DNA not treated with bisulfite was amplified as a loading control. The PCR products were then electrophoresed on 3 % agarose gel and visualized under UV illumination using the Fusion FX Imaging system (Vilber). The ratio of methylated to non-methylated DNA was calculated after gel intensity quantification in Fiji / ImageJ.

### Immunofluorescent staining

Adrenal glands cleaned from surrounding fat tissue were fixed in 4 % PFA in PBS, washed overnight in PBS, cryopreserved in 30 % sucrose (AppliChem GmbH) in PBS overnight at 4°C, embedded in O.C.T. Compound (Tissue-Tek) and frozen at −80 °C. Each adrenal gland was cut into 8 μm thick serial sections. Before staining, adrenal sections were pre-warmed at RT for 30 min and antigen retrieval was performed by boiling in citrate buffer (pH 6) for 6 min. Adrenal sections were washed with PBS, permeabilized with 0.1 % Triton X-100 in PBS for 20 min, treated with TrueBlack Lipofuscin Quencher (1:40 in 70 % ethanol; Biotium) for 30 sec to reduce autofluorescence and blocked in Dako Protein Block, serum-free buffer for 1 hour at RT. Then, sections were incubated overnight at 4 °C with primary antibodies, washed with PBS, and incubated for 1 hour at RT with the secondary antibodies together with DAPI (1:5,000; Roche), all diluted in Dako Antibody Diluent. Antibodies and dyes used were: anti-SDHB (1:300; Sigma-Aldrich, #HPA002868), anti-IDH2 (1:50; Sigma-Aldrich, #HPA007831), anti-SF-1 (1:100; TransGenic Inc. #KO610), Lectin Esculentum DyLight488 (1:300; Vector Laboratories, #DL-1174), 4-Hydroxynonenal (4-HNE, 1:200; Abcam, # ab48506), Alexa Fluor 555 donkey anti-rabbit (1:300; Life Technologies, #A-31572), Alexa Fluor 647 chicken anti-rat (1:300; Invitrogen, #A21472) and Alexa Fluor 555 donkey anti-mouse (1:300; Invitrogen, #A31570). After washing with PBS, cryosections were mounted with Fluoromount (Sigma-Aldrich), covered with 0.17 mm cover glass, fixed with nail polish and kept at 4 °C until imaging.

### Image acquisition and image analysis

Z-series microscopic images for SDHB and IDH2 staining were acquired on Zeiss LSM 880 inverted confocal microscope (Zeiss, Jena, Germany), illuminated with laser lines at 405 nm, 488 nm, 561 nm and 633 nm, and detected by two photomultiplier tube detectors. EC Plan-Neofluoar objective with 40x magnification, 1.30 numerical aperture and M27 thread, working with an oil immersion medium Immersol 518 F, was used. Microscopic images of SF-1 and 4-HNE stainings were acquired with an Axio Observer Z1/7 inverted microscope with Apotome mode (Zeiss, Jena, Germany), illuminated with LED-Module 385 nm and 567 nm, on a Plan-Apochromat objective with 10x magnification, 0.45 numerical aperture and M27 thread. Laser power, photomultiplier gain and pinhole size were set for each antibody individually and kept constant for all image acquisitions. For each condition, at least 3 view-fields were imaged per tissue section. Images were acquired with the ZEN 3.2 blue edition software, and processed and quantified with the Fiji / ImageJ software on maximum intensity Z-projection images.

### Statistical analysis

The statistical analysis and data plotting were done with the GraphPrism 7 software. The statistical tests used are described in each figure legend, p < 0.05 was set as a significance level.

### Graphical design

Figure 7 was created with Biorender.com.

**Fig. 1.**
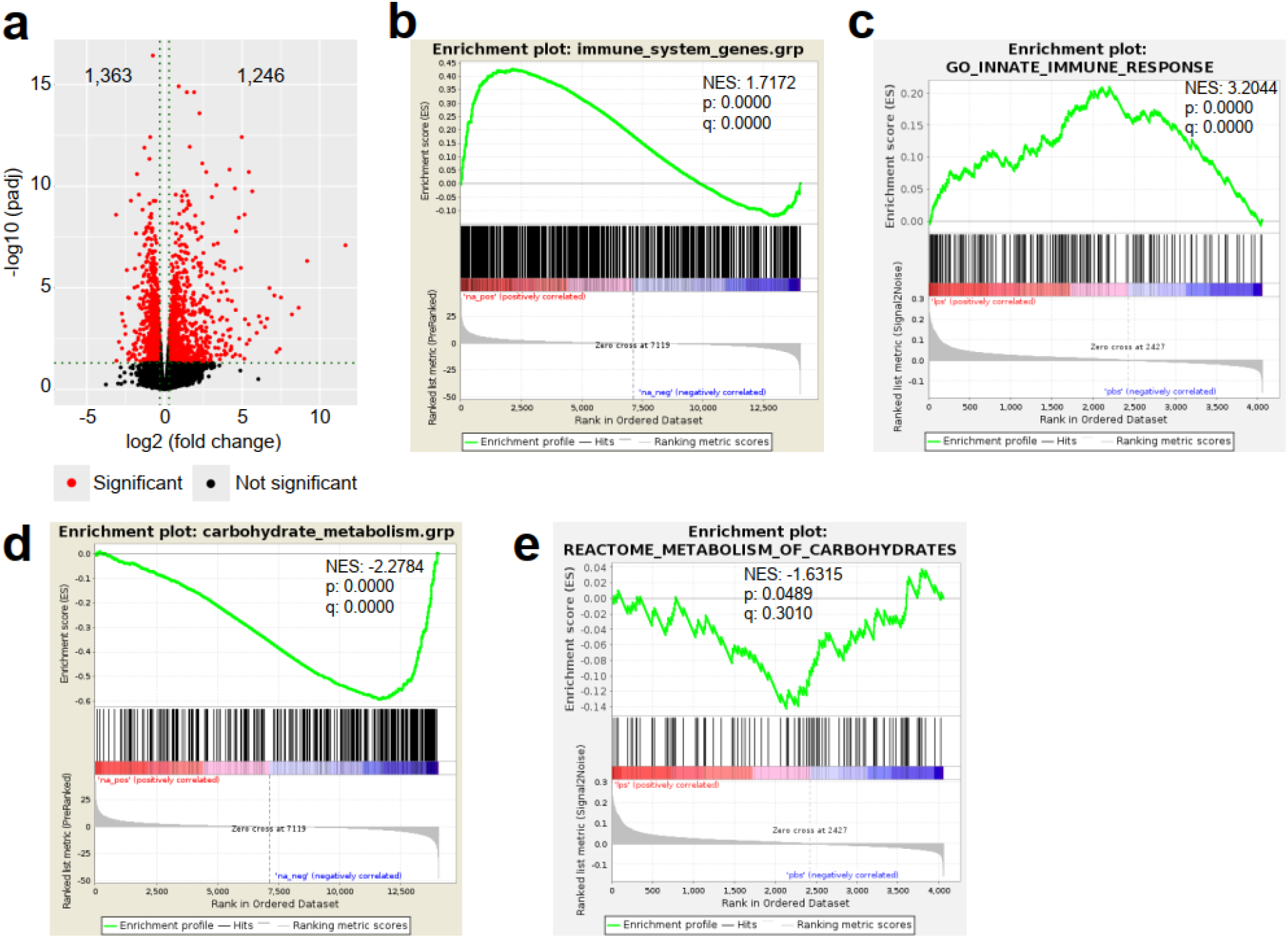
LPS-induced inflammation changes the transcriptional and proteomic profile of the adrenal cortex. **a** Volcano plot showing differentially expressed genes in the microdissected adrenal gland cortex of mice treated for 6 h with PBS or LPS. **b** GSEA for immune pathways in the adrenal cortex of LPS-versus PBS mice. **c** GSEA for proteins associated with the innate immune response in CD31^−^CD45^−^ adrenocortical cells of mice treated for 24 h with PBS or LPS. **d** RNA-Seq based GSEA for carbohydrate metabolism in the adrenal cortex of LPS-versus PBS mice. **e** GSEA for proteins associated with carbohydrate metabolism in CD31^−^CD45^−^ adrenocortical cells of LPS versus PBS mice. NES: normalized enrichment score. **a**,**b**,**d** n = 3 mice per group, **c,e** n = 6 mice per group, padj < 0.05 was used as a cut-off for significance.

**Fig. 2.**
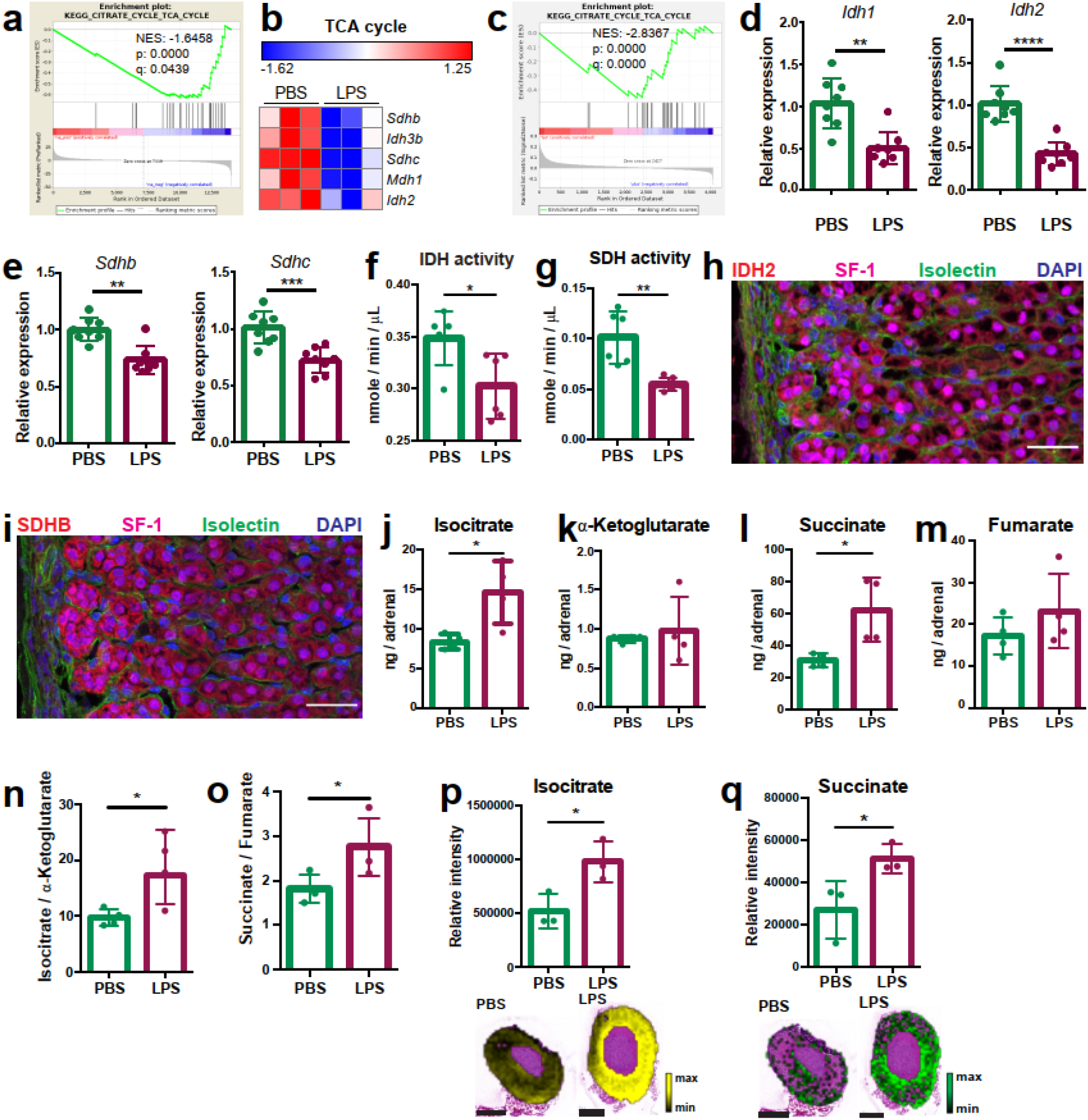
Systemic inflammation disrupts the TCA cycle in the adrenal cortex. **a,b** Transcriptome analysis in the microdissected adrenal gland cortex of mice treated for 6 h with PBS or LPS (n = 3 mice per group). **a** GSEA for TCA cycle genes. **b** Heatmap of differentially expressed TCA cycle genes (padj < 0.05). **c** GSEA analysis for TCA cycle proteins in CD31^−^CD45^−^ adrenocortical cells of mice treated for 24 h with PBS or LPS (n = 6 mice per group). **d**,**e** mRNA expression of *Idh1, Idh2, Sdhb* and *Sdhc* in adrenocortical CD31^−^CD45^−^ cells of mice treated for 6 h with PBS or LPS (n = 8 mice per group, shown is one from two experiments). **f,g** Quantification of IDH and SDH activities in the adrenal cortex of mice treated for 24 h with LPS or PBS (n = 6 mice per group). Values are normalized to the total protein amount in the adrenal cortex. **h**,**i** Immunofluorescence images of the adrenal gland, stained for IDH2 (red) or SDHB (red), SF-1 (magenta), Isolectin (staining endothelial cells, green), and DAPI (blue). Scale bar, 30 μm. **j-o** TCA cycle metabolites (isocitrate, α-ketoglutarate, succinate, fumarate) were measured by LC–MS/MS in adrenal glands of mice 24 h after injection with PBS or LPS (n = 4 mice per group, shown is one from two experiments). **p,q** MALDI-MSI for isocitrate and succinate in the adrenal cortex of mice treated for 24 h with PBS or LPS (n = 3 mice per group). Representative images and quantifications are shown. Scale bar, 500 μm. Data are presented as mean ± s.d., and analyzed with Mann-Whitney test. *p < 0.05, **p < 0.01, ***p < 0.001, ****p < 0.0001.

**Fig. 3.**
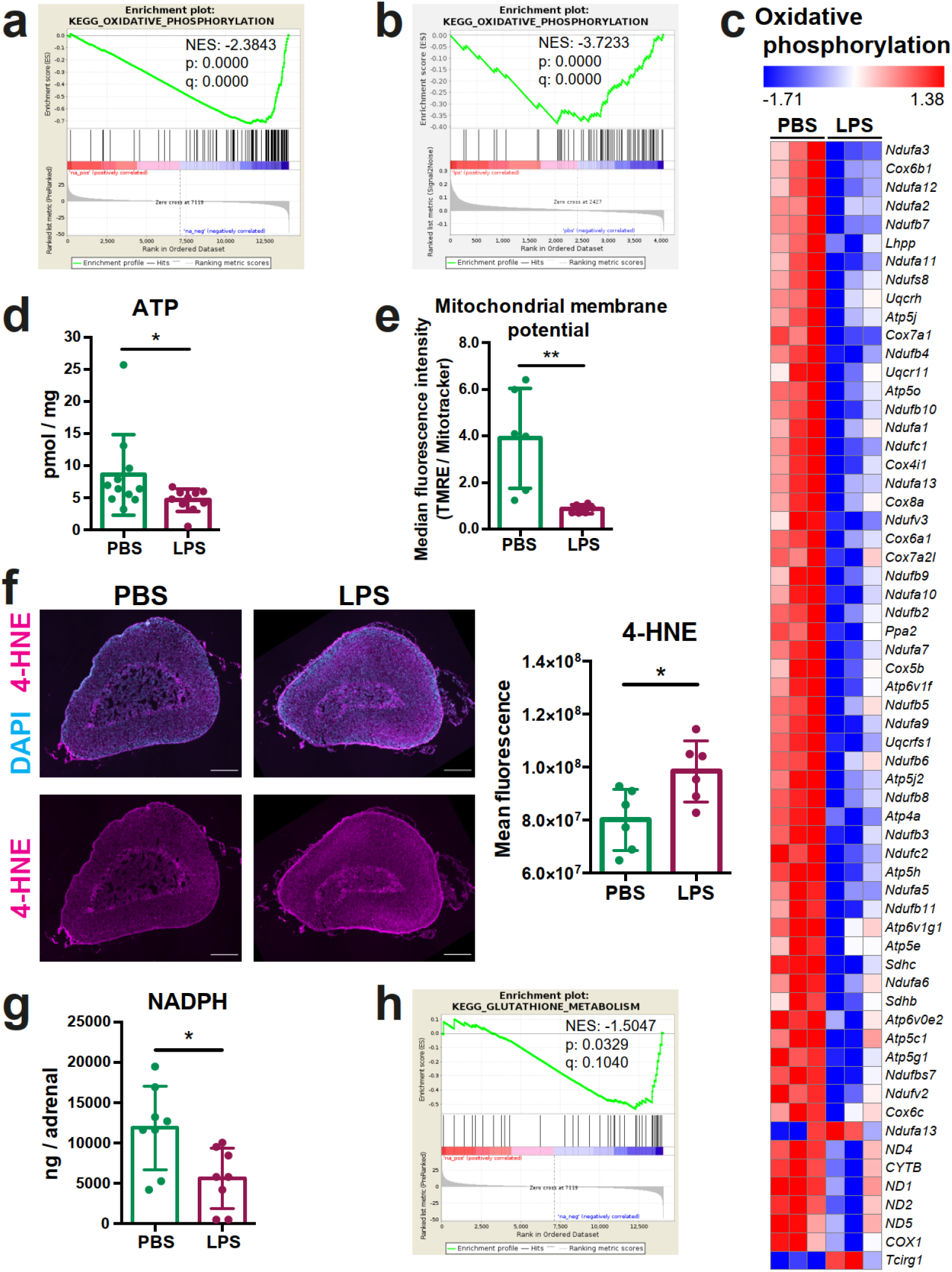
Oxidative phosphorylation is reduced and oxidative stress is increased in the adrenal cortex of LPS treated mice. **a** GSEA for oxidative phosphorylation-related genes in the adrenal cortex of mice treated for 6 h with PBS or LPS (n = 3 mice per group). **b** GSEA for oxidative phosphorylation-associated proteins in CD31^−^CD45^−^ adrenocortical cells of mice treated for 24 h with PBS or LPS (n = 6 mice per group). **c** Heatmap of differentially expressed genes related to oxidative phosphorylation (padj < 0.05). **d** Measurement of ATP in adrenal glands of mice treated for 24 h with PBS or LPS (n = 10-11 mice per group). **e** Measurement of mitochondrial membrane potential by TMRE staining and mitochondrial load by Mitotracker Green FM in CD31^−^CD45^−^adrenocortical cells of PBS or LPS mice. Data are presented as ratio of the median fluorescence intensities of TMRE to Mitotracker Green FM (n = 6 mice per group). **f** Representative immunofluorescence images of adrenal gland sections from PBS- and LPS-treated mice (24 h post-injection), stained for 4-HNE (magenta) and DAPI (blue). Scale bar, 300 μm. Quantification of the mean fluorescence intensity of 4-HNE staining in the adrenal cortex of PBS- or LPS-treated mice (n = 6 mice per group). **g** NADPH measurement by LC–MS/MS in adrenal glands of mice treated with PBS or LPS for 24 h (n = 8 mice per group). Data are given as observed peak area intensities of NADPH. **h** GSEA for glutathione metabolism of RNA-Seq data in the adrenal cortex of LPS versus PBS mice. Data present mean ± s.d. and were analyzed with Mann-Whitney test. *p < 0.05, **p < 0.01.

**Fig. 4.**
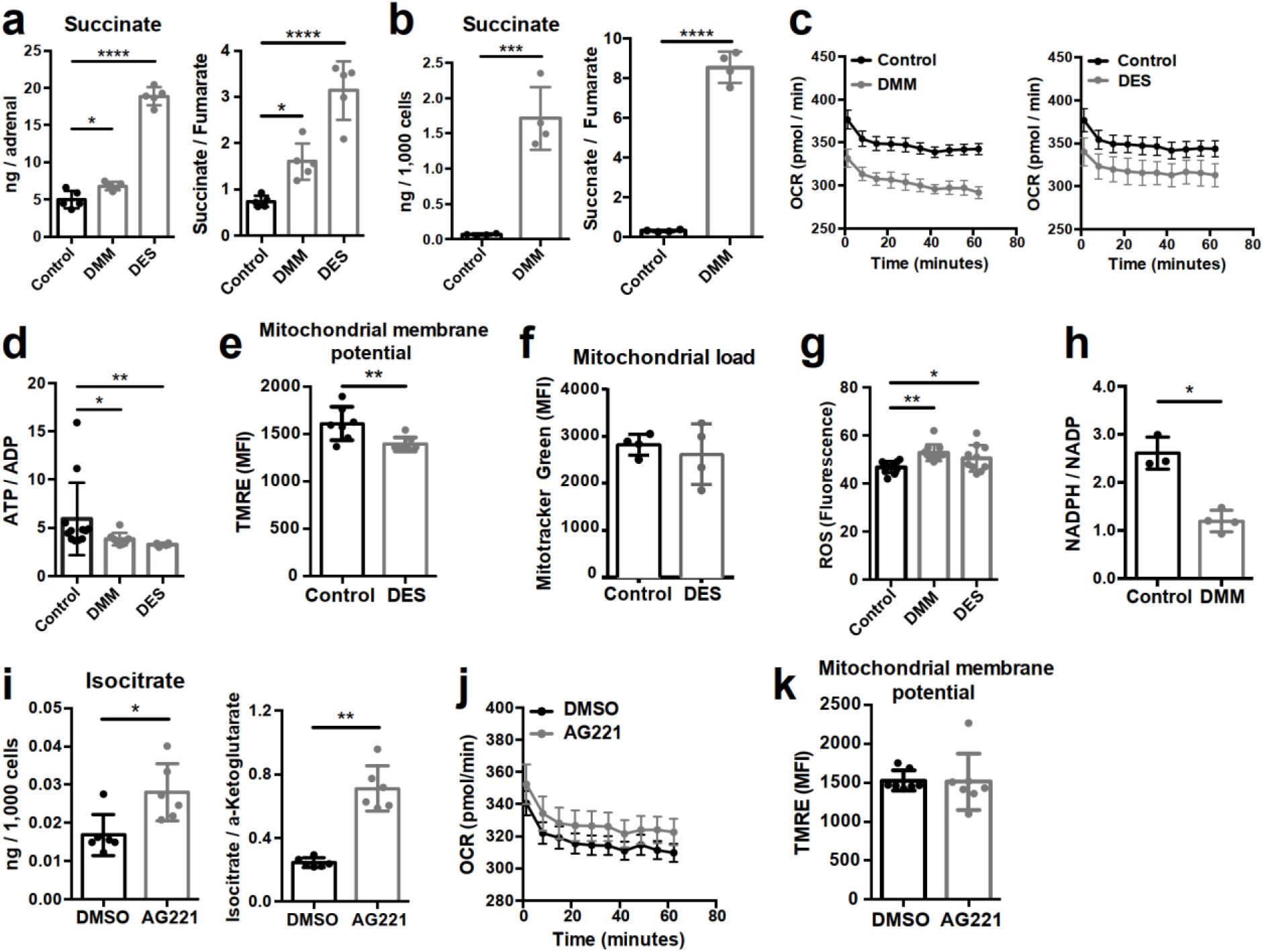
Increased succinate levels impair mitochondrial function in adrenocortical cells. **a**,**b** Succinate and fumarate levels were measured by LC–MS/MS in adrenal gland explants (**a**) and NCI-H295R cells (**b**) treated with DMM or DES for 24 h (n = 5 and n = 4 for **a** and **b**, respectively). **c** OCR measurement with Seahorse technology in NCI-H295R cells treated with DMM or DES for 24 h (n = 6). **d** Measurement of ATP/ADP ratio in NCI-H295R cells treated with DMM or DES for 24 h (n = 4-8). **e**,**f** TMRE and Mitotracker Green FM staining assessed by flow cytometry in NCI-H295R cells treated with DES for 4 h, MFI is shown (n = 7 for **e** and n = 4, one from two experiments for **f**). **g** ROS measurement in NCI-H295R cells treated with DMM or DES for 2 h (n = 10 - 12). **h** Measurement of NADPH/NADP ratio in NCI-H295R cells treated with DMM for 24 h (n = 3 - 4). **i** Isocitrate levels measured by LC–MS/MS in NCI-H295R cells treated for 24 h with AG221 or DMSO (n = 6). **j** OCR measurement in NCI-H295R cells treated for 24 h with AG221 or DMSO (n = 10). **k** TMRE staining and flow cytometry in NCI-H295R cells treated for 4 h with AG221 or DMSO, MFI is shown (n = 7). Data are presented as mean ± s.d., and analyzed with one-way ANOVA (**a**,**d**,**g**) or Mann-Whitney test (**b**,**e**,**f**,**h**,**i**,**k**). Data in **c** and **j** are shown as mean ± s.e.m. *p < 0.05, **p < 0.01, ***p < 0.001, ****p < 0.0001.

**Fig. 5.**
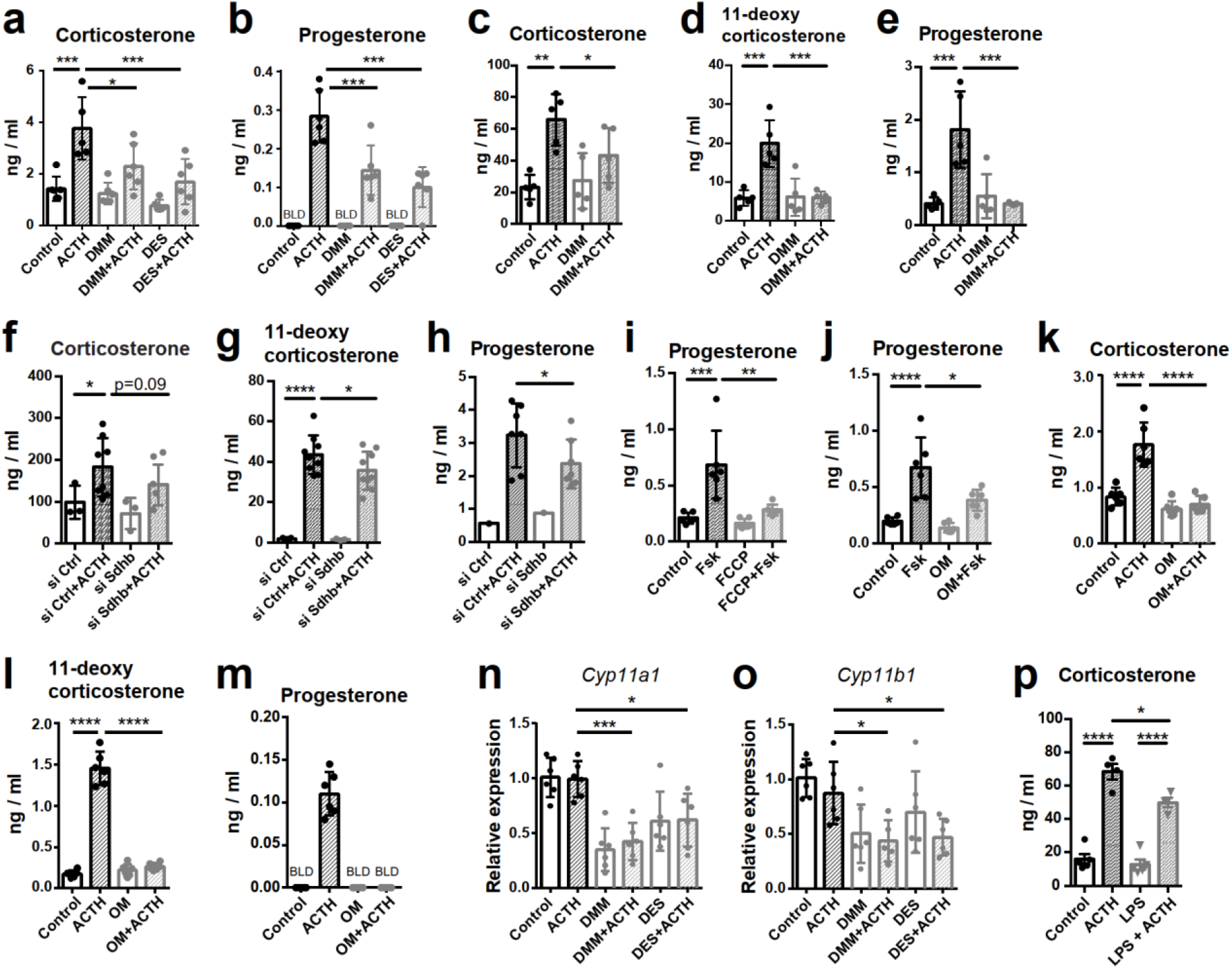
Disruption of SDH function impairs glucocorticoid production. **a**-**e** Primary adrenocortical cells (**a**,**b**) and adrenal explants (**c-e**) were treated for 24 h with DMM or DES and for another 45 min with ACTH (10 ng/mL or 100 ng/mL, respectively) (n = 5-6). **f-g** Primary adrenocortical cells were transfected with *siSdhb* or non-targeting siRNA (siCtrl) and 24 h post-transfection they were treated for 45 min with ACTH (n = 7-8). **i**,**j** NCI-H295R cells were treated for 24 h with FCCP (**i**) or oligomycin (OM) (**j**) and for another 30 min with Forskolin (Fsk) (n = 6). **k-m** Primary adrenocortical cells were treated for 24 h with oligomycin (OM) and for another 45 min with ACTH (n = 6). **n,o** *Cyp11a1* and *Cyp11b1* expression in primary adrenocortical cells treated for 24 h with DMM or DES and for 45 min with ACTH (n = 5-6). **p** Adrenal gland explants were treated for 24 h with LPS and for 45 min with ACTH (n = 4-5). Measurements of steroid hormones in **a-m,p** were performed in supernatants of primary adrenocortical cell cultures or adrenal gland explants by LC-MS/MS. Data are shown as mean ± s.d. and analyzed with one-way ANOVA. *p < 0.05, **p < 0.01, ***p < 0.001, ****p < 0.0001. BLD = below level of detection.

**Fig. 6.**
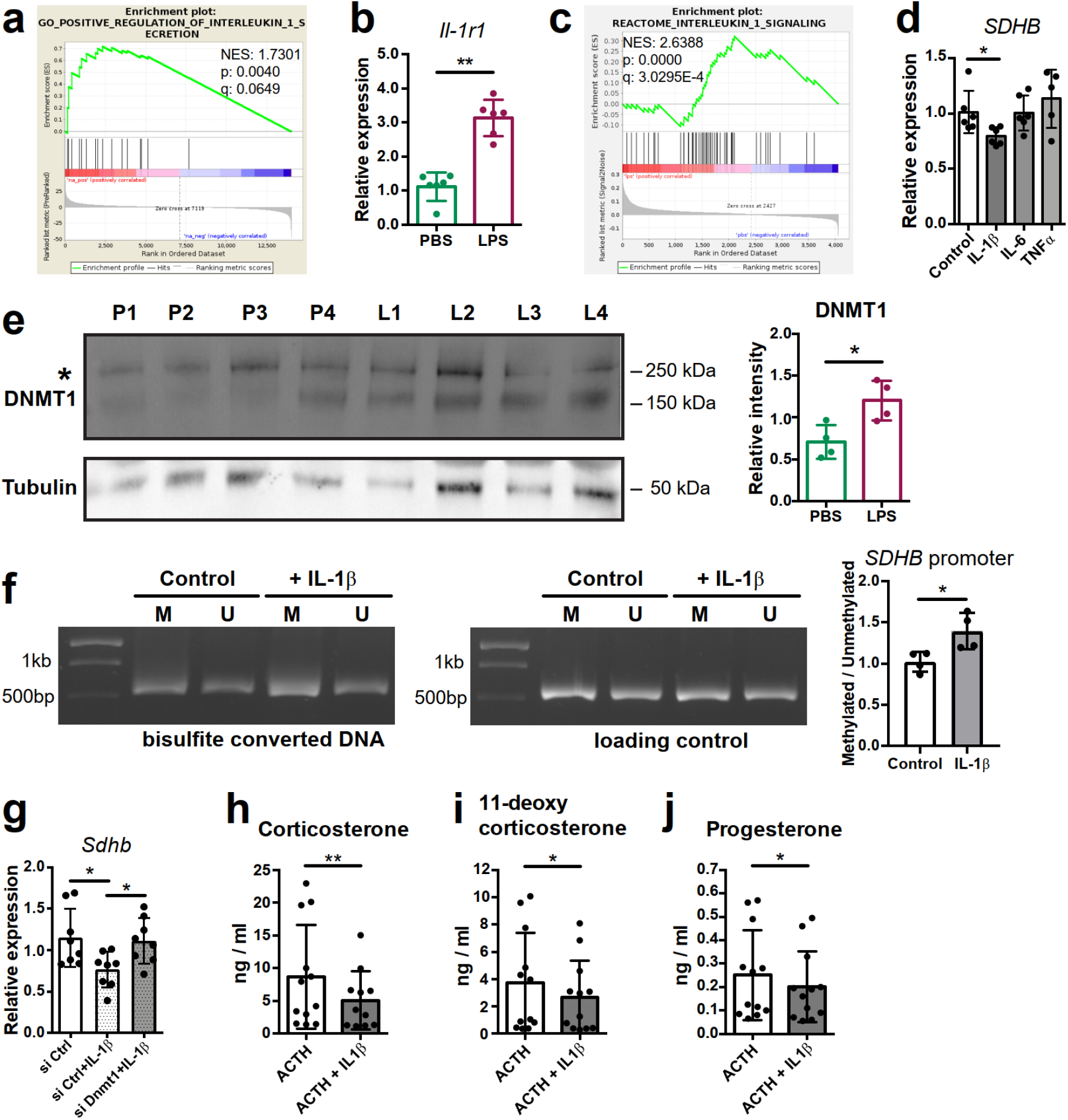
IL-1β reduces SDHB expression and glucocorticoid synthesis in a DNMT1-dependent manner. **a** GSEA for genes related to positive regulation of IL-1β secretion in the adrenal cortex of mice treated for 6 h with PBS or LPS (n = 3 mice per group). **b** *Il-1r1* expression in CD31^−^CD45^−^ adrenocortical cells of mice 6 h post-injection with PBS or LPS (n = 6 mice per group). **c** GSEA for proteins related to IL-1 β signaling in CD31^−^CD45^−^ adrenocortical cells of mice treated for 24 h with PBS or LPS (n = 6 mice per group). **d** *SDHB* expression in NCI-H295R cells treated for 2 h with IL-1β, IL-6 or TNFα (n = 6). **e** Western blot analysis for DNMT1 in CD31^−^CD45^−^ adrenocortical cells of mice 24 h after injection of PBS (P) or LPS (L) (n = 4 mice per group), α-TUBULIN was used as loading control. The asterisk (*) depicts an unspecific band. Quantification of the western blot, shown as relative intensity of DNMT1 to α-TUBULIN. **f** NCI-H295R cells were treated for 2 h with IL-1β (n = 4), and the ratio of methylated to unmethylated DNA in the *SDHB* promoter was assayed after bisulfite conversion. Representative gel electrophoresis images of bisulfite converted and non-treated DNA (M – methylated, U – unmethylated). **g** *Sdhb* expression in primary adrenocortical cells transfected with *siDnmt1* or siCtrl and 24 h post-transfection treated for 6 h with IL-1β (n = 7-8). **h-j** Primary adrenocortical cells were treated for 6 h with IL-1β and for another 45 min with ACTH (10 ng/mL) (n = 11-12). Steroid hormones were measured in the culture supernatant by LC-MS/MS. Data are shown as mean ± s.d. and analyzed with Mann-Whitney (**b**,**d**,**e**,**f**) or Wilcoxon test (**g-j**). *p < 0.05, **p < 0.01.

**Fig. 7.**
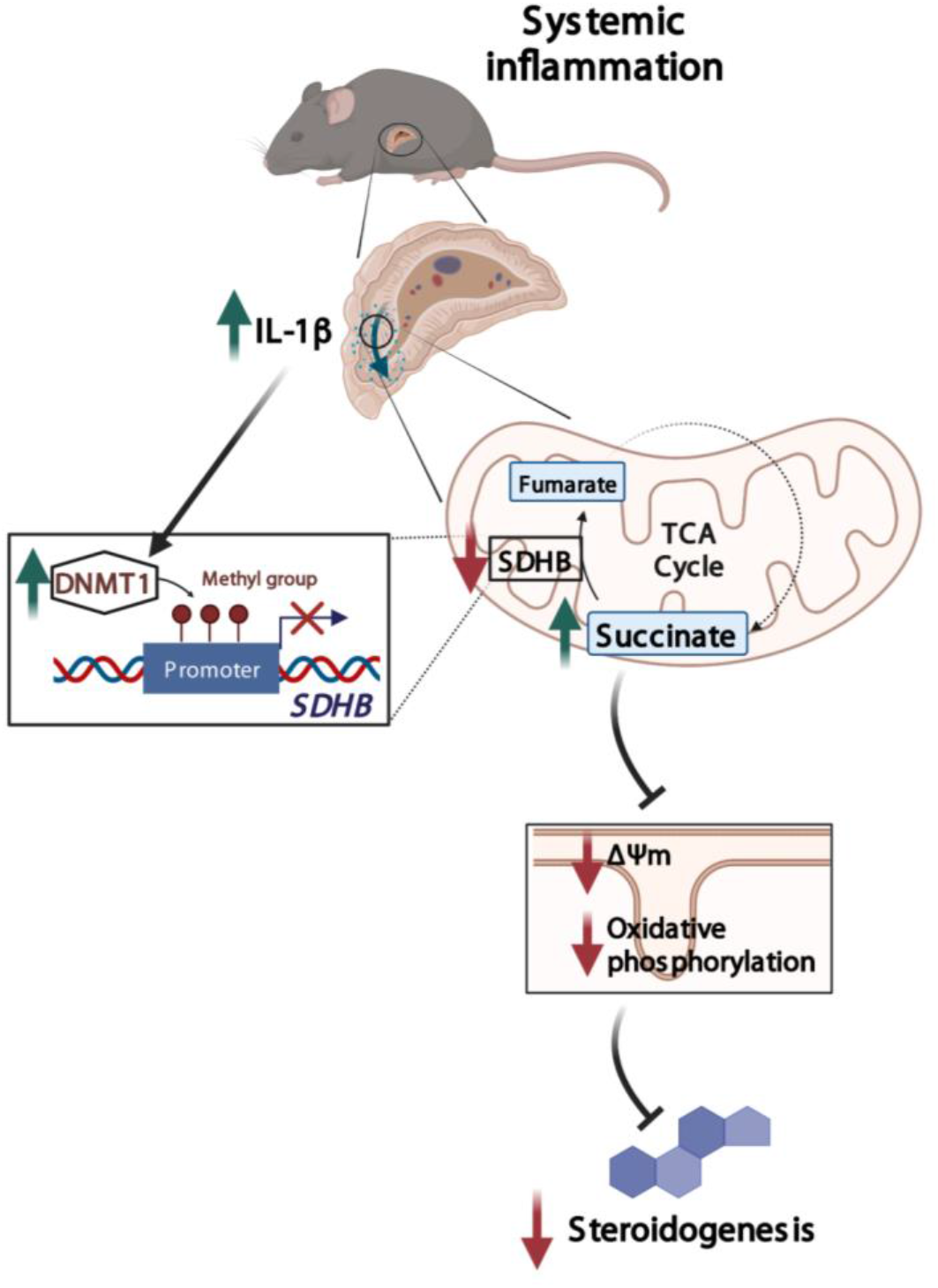
Illustration of the regulation of adrenocortical steroidogenesis by inflammation. IL-1β reduces SDHB expression through upregulation of DNA methyltransferase 1 (DNMT1) and methylation of the SDHB promoter. Consequently, increased succinate levels impair oxidative phosphorylation, leading to reduced steroidogenesis.

## Results

### 1. Metabolic reprograming of the adrenal cortex in inflammation

To explore inflammation-induced alterations in the adrenal cortex, we performed RNA-Seq in microdissected adrenal cortices from mice treated for 6 h i.p. with 1 mg / kg LPS or PBS, which revealed 2,609 differentially expressed genes, out of which 1,363 were down- and 1,246 were upregulated (Fig. 1a). Gene set enrichment analysis (GSEA) using the Molecular Signatures Database (MSigDB) hallmark gene set collection [39] showed a significant enrichment of inflammatory response-related gene sets in the adrenal cortex of LPS-treated mice (Fig. 1b). In acute inflammation, leukocytes infiltrate the adrenal cortex [7, 9] and resident macrophages are activated [40,41]. In order to delineate the inflammatory response in the adrenocortical steroidogenic cells, CD31^−^CD45^−^ cells were sorted: enrichment in steroidogenic cells was evidenced by high *Star* expression (Suppl. Fig. 1a), and purity was verified by the absence of *Cd31* and *Cd45* expression (Suppl. Fig. 1b,c). Proteomic analysis in the sorted CD31^−^CD45^−^ adrenocortical cell population and GSEA of GO terms confirmed the enrichments of innate immune response-related proteins in adrenocortical cells of LPS-injected mice (Fig. 1c), suggesting that steroidogenic adrenocortical cells respond to inflammatory stimuli.

LPS treatment leads to increased plasma corticosterone levels [7, 9, 10]. Numerous studies have shown that elevated glucocorticoid levels are primarily driven by activation of the HPA axis, and coincide with increased circulating ACTH levels [7, 9, 10]. This is accompanied by increased expression of genes related to steroid biosynthesis [42]. We confirmed an increased expression of the cholesterol transporter *Star* [43] and the terminal enzyme for glucocorticoids synthesis *Cyp11b1* [2] in adrenocortical cells of LPS mice (Suppl. Fig. 2a,b). However, the expression of other steroidogenic enzymes, such as *3βHSD* and *Cyp21a1* was reduced, while *Cyp11a1* remained unchanged (Suppl. Fig. 2c-e). Similarly, protein levels of the transcription factor SF-1, a key inducer of steroidogenesis [44], were somewhat reduced after LPS injection (Suppl. Fig. 2f). Therefore, the observed changes in plasma glucocorticoid levels which accompany inflammation cannot be solely explained by the transcriptional changes in steroidogenic enzymes.

Next, we explored the cell metabolic changes induced by LPS in the adrenal cortex. By GSEA of the RNA-Seq data we observed negative regulation of gene sets related to carbohydrate metabolism in the adrenal cortex of LPS-injected mice (Fig. 1d). Proteomic analysis was performed in CD31^−^CD45^−^ adrenocortical cells (Suppl. Fig. 1) to examine the effects of inflammation on the metabolism of specifically steroidogenic adrenocortical cells, thus evading the well-described inflammation-induced metabolic changes in immune cells [16–18]. Similarly to the RNA-Seq data, GSEA of the proteomic data showed significant negative enrichment of proteins associated with carbohydrate metabolism in the steroidogenic cells (Fig. 1e). EGSEA pathway analysis of the RNA-Seq and proteomic data revealed that TCA cycle, oxidative phosphorylation, tyrosine metabolism, fatty acid degradation, D-Glutamine and D-glutamate metabolism, glutathione metabolism and other metabolic pathways were significantly enriched among the downregulated genes and proteins in the adrenal cortex and in steroidogenic cells of LPS mice (Table 1, Table 2).

### 2. Inflammation disrupts the TCA cycle in adrenocortical cells at the levels of isocitrate dehydrogenase and succinate dehydrogenase

Inflammation downregulates the TCA cycle and oxidative phosphorylation in inflammatory activated macrophages [16], however little is known about inflammation-induced metabolic changes in other cell types. We show that TCA cycle-related gene expression was downregulated in the adrenal cortex of LPS-treated mice (Fig. 2a,b, Table 1). Expression of genes encoding key TCA cycle enzymes, including succinate dehydrogenase *Sdhb* and *Sdhc*, isocitrate dehydrogenase 2 and 3 (*Idh2* and *Idh3b*) and malate dehydrogenase 1 (*Mdh1*), was reduced in the adrenal cortex of LPS-injected mice (Fig. 2b). Proteomic GSEA confirmed the TCA cycle downregulation in steroidogenic adrenocortical cells of LPS mice (Fig. 2c, Table 2). Accordingly, CD31^−^CD45^−^ adrenocortical cells from LPS mice displayed reduced *Idh1, Idh2, Sdhb* and *Sdhc* expression (Fig. 2d,e) and LPS treatment attenuated the IDH and SDH enzymatic activities in the adrenal cortex (Fig. 2f,g). Additionally, immunofluorescent staining showed that IDH2 and SDHB proteins are highly expressed in SF-1^+^ (steroidogenic) cells (Fig. 2h,i). In endothelial and immune cells of the adrenal cortex of LPS mice, *Idh1* and *Idh2* gene expression was reduced, *Sdhb* gene expression was increased, while expression of *Sdhc* was unaltered (Suppl. Fig. 3a,b). Collectively, these data indicate that the reduced activity of SDH and IDH in the adrenal cortex of LPS-treated mice is mainly due to their downregulated expression in steroidogenic adrenocortical cells.

In order to confirm that inflammation disrupts the TCA cycle in adrenocortical cells, we profiled the changes in metabolite levels in the adrenal glands of PBS- and LPS-treated mice using liquid chromatography – tandem mass spectrometry (LC-MS/MS). The levels of isocitrate and succinate, as well as the ratios of isocitrate / α-ketoglutarate and succinate / fumarate were increased in the adrenal glands of LPS-treated mice (Fig. 2j-o). Furthermore, MALDI MS–Imaging confirmed the increased levels of isocitrate and succinate in the adrenal cortex of LPS mice (Fig. 2p,q). These data collectively demonstrate that inflammation disrupts IDH and SDH activities and increases the levels of their substrates isocitrate and succinate in adrenocortical cells.

### 3. Inflammation reduces oxidative phosphorylation and increases oxidative stress in the adrenal cortex

Next, we investigated how inflammation affects mitochondrial oxidative metabolism in adrenocortical cells. GSEA of the RNA-Seq and proteomic data in the adrenal cortex and CD31^−^CD45^−^ adrenocortical cells, respectively, revealed that oxidative phosphorylation was significantly enriched among the downregulated genes (Fig. 3a) and proteins (Fig. 3b), and expression of a large number of oxidative phosphorylation-associated genes was reduced in the adrenal cortex of LPS mice (Fig. 3c). In accordance, ATP levels were reduced in the adrenal gland (Fig. 3d) and the mitochondrial membrane potential of CD31^−^CD45^−^ adrenocortical cells was decreased in mice treated with LPS (Fig. 3e). In pro-inflammatory macrophages, a TCA cycle ‘break’ at the level of SDH is associated with repurposing of mitochondria from oxidative phosphorylation-mediated ATP synthesis to ROS production [45]. EGSEA pathway analysis showed that upon LPS treatment several pathways involved in the regulation of and the cellular response to oxidative stress in the adrenal cortex were enriched at mRNA (Table 3) and protein level (Table 4). This was confirmed by increased 4-hydroxynonenal (4-HNE) staining, indicating higher oxidative stress-associated damage in the adrenal cortex of LPS-treated mice (Fig. 3f). Antioxidant defense mechanisms are particularly important in the adrenal cortex, since electron leakage through the reactions catalyzed by CYP11A1 and CYP11B1 during glucocorticoid synthesis contributes significantly to mitochondrial ROS production [46]. Cells neutralize ROS to maintain their cellular redox environment by using the reducing equivalents NADPH and glutathione [47]. In addition, NADPH serves as a co-factor for mitochondrial steroidogenic enzymes [48]. NADPH levels and glutathione metabolism-related gene expression were significantly decreased in the adrenal glands of LPS mice (Fig. 3g,h, Table 1, Table 2). These findings collectively suggest that inflammation in the adrenal cortex is associated with increased oxidative stress, perturbed mitochondrial oxidative metabolism, reduced anti-oxidant capacity and increased ROS production.

### 4. Increased succinate levels impair mitochondrial metabolism and steroidogenesis in adrenocortical cells

SDH is complex II of the electron transport chain (ETC), coupling succinate oxidation with the respiratory chain [49]. Inhibition of SDH function with dimethyl malonate (DMM), which is hydrolyzed to the competitive SDH inhibitor malonate [24/50], or treatment of adrenocortical cells with the cell-permeable succinate analog diethyl succinate (DES) increased the amount of succinate and the succinate / fumarate ratio in adrenal gland explants (Fig. 4a) and human adrenocortical carcinoma cells NCI-H295R (Fig. 4b) and decreased the oxygen consumption rate (OCR) and ATP production in the latter (Fig. 4c,d). This was associated with reduced mitochondrial membrane potential (Fig. 4e), but not mitochondrial load (Fig. 4f). Furthermore, DMM and DES increased ROS (Fig. 4g) and decreased NADPH levels (Fig. 4h), suggesting that in adrenocortical cells, as in macrophages [45], succinate repurposes mitochondrial metabolism from oxidative phosphorylation to ROS production. Such changes in the mitochondrial function were not observed when inhibiting IDH activity with enasidenib (AG221) [51] (Fig. 4i-k). AG221 increased isocitrate and the isocitrate / α-ketoglutarate ratio (Fig. 4i), but did not affect OCR (Fig. 4j) or the mitochondrial membrane potential (Fig. 4k).

Key steps of steroidogenesis take place in the mitochondria [49], thus, we asked whether disruption of SDH activity affects steroidogenic function. We inhibited SDH activity with DMM in human and mouse adrenocortical cells, and induced glucocorticoid production by forskolin or ACTH, respectively. SDH inhibition considerably impaired glucocorticoid and progesterone production in mouse primary adrenocortical cells (Fig. 5a,b), adrenal gland explants (Fig. 5c-e) and human adrenocortical NCI-H295R cells (Suppl. Fig. 4a,b). Similarly, DES diminished glucocorticoid production in mouse (Fig. 5a,b) and human adrenocortical cells (Suppl. Fig. a,b). Confirming these data, *Sdhb* silencing (Suppl. Fig. 5a,b) impaired glucocorticoid synthesis in mouse (Fig. 5f-h) and human adrenocortical cells (Suppl. Fig. 4c), implying that proper adrenocortical steroidogenesis relies on intact SDH activity. Recently it was shown that SDH activity and intracellular succinate are required for CYP11A1-mediated pregnenolone synthesis, the first step of steroidogenesis [52]. Adding to this knowledge, our data demonstrate that increasing succinate concentrations impair steroidogenesis (Suppl. Fig. 4d-f). Moreover, the proton gradient uncoupler FCCP (Fig. 5i) and the ATP synthase inhibitor oligomycin (Fig. 5j-m) both strongly reduced steroidogenesis in mouse (Fig. 5k-m) and human adrenocortical cells (Fig. 5i,j), demonstrating the well-established requirement of intact mitochondrial membrane potential and ATP generation for steroidogenic function [52,53]. In accordance, DMM and DES downregulated the expression of *Cyp11a1* and *Cyp11b1* (Fig. 5n,o), genes encoding for enzymes with critical roles in glucocorticoid synthesis, that catalyze the conversion of cholesterol to pregnenolone and the final step of corticosterone/cortisol production, respectively [2, 49]. Importantly, treatment of adrenal gland explants with LPS reduced corticosterone secretion in response to ACTH, similarly to DMM of DES (Fig. 5p).

In contrast, inhibition of IDH activity with AG221 (Fig. 4i), did not alter glucocorticoid production in mouse adrenocortical cells (Suppl. Fig. 6a,b), adrenal gland explants (Suppl. Fig. 6c,d) or human adrenocortical cells (Suppl. Fig. 6e,f), nor did *Idh2* silencing in mouse adrenocortical cells (Suppl. Fig. 5c, 6g,h). Taken together, these results imply that SDH but not IDH activity is required for adrenocortical steroidogenesis.

### 5. Itaconate is not responsible for reduced SDH activity and steroidogenesis in adrenocortical cells

In inflammatory macrophages SDH function is inhibited by itaconate [54], a byproduct of the TCA cycle produced from cis-aconitate in a reaction catalyzed by aconitate decarboxylase 1 (ACOD1) [55]. The expression of *Irg1*, the gene encoding for ACOD1, and itaconate levels are strongly upregulated in macrophages upon inflammation [54]. We asked whether itaconate might affect SDH activity in the adrenal cortex. *Irg1* expression was upregulated in the adrenal cortex of LPS-treated mice but this increase derived from CD45^+^ cells, while *Irg1* was not expressed in CD31^−^CD45^−^ adrenocortical cells (Suppl. Fig. 7a). Accordingly, LPS treatment significantly elevated itaconate levels in the CD31^+^CD45^+^ fraction, while it did not increase itaconate levels in CD31^−^CD45^−^ adrenocortical cells (Suppl. Fig. 7b,c). Itaconate can be secreted from LPS-stimulated macrophages [54], and could thereby affect SDH activity in adrenocortical cells. Therefore, we tested whether exogenously given itaconate may affect steroidogenesis by treating primary adrenocortical cells with the cell permeable itaconate derivative 4-octyl itaconate (4-OI). Adrenocortical cells internalized the added itaconate derivative (Suppl. Fig. 7d), which however did not alter succinate or fumarate levels or the succinate / fumarate ratio (Suppl. Fig. 7e-g), nor did it affect glucocorticoid production (Suppl. Fig. 7h,i). Additionally, SDH activity in the adrenal cortex of *Irg1-KO* mice injected with LPS was not different from that in their wild-type counterparts (Suppl. Fig. 7j). Hence, neither is itaconate produced nor does it affect SDH activity through paracrine routes in adrenocortical cells.

### 6. IL-1β downregulates SDHB expression and steroidogenesis in a DNMT1-dependent manner

Systemic inflammation induces substantial leukocyte recruitment in the adrenal gland, accompanied by elevated production of pro-inflammatory cytokines [7, 56]. IL-1β is highly produced by inflammatory monocytes and macrophages [57]. RNA-Seq in the adrenal cortex, including recruited immune cells, showed increased expression of *Il-1β* in LPS- compared to PBS-injected mice (log2fc = 1.46, padj = 0.019). Furthermore, there was significant positive enrichment of genes associated with IL-1β secretion in the adrenal cortex of mice treated with LPS (Fig. 6a). The IL-1β receptor *Il-1r1* is expressed in CD31^−^CD45^−^ adrenocortical cells and its expression was upregulated in adrenocortical cells sorted from LPS treated mice (Fig. 6b). In accordance, proteins related to IL-1β signaling were positively enriched in CD31^−^CD45^−^ adrenocortical cells of LPS mice (Fig. 6c). Essentially, IL-1β, but not IL-6 or TNFα, reduced *SDHB* expression in NCI-H295R cells (Fig. 6d). One way of transcriptional gene repression is covalent attachment of methyl groups on the cytosine 5’ position within the gene promoter sequence, a reaction catalyzed by DNA methyltransferases [58]. Proteomics revealed significant upregulation of DNA methyltransferase 1 (DNMT1) in CD31^−^CD45^−^ adrenocortical cells of LPS mice (log2fc = 0.421, padj = 0.015), which we confirmed by western blot analysis (Fig. 6e). In accordance, IL-1β increased DNA methylation of the *SDHB* promoter (Fig. 6f), and *Dnmt1* silencing restored *Sdhb* expression in IL-1β treated adrenocortical cells (Fig. 6g, Suppl. Fig. 5d). Lastly, IL-1β reduced adrenocortical steroidogenesis (Fig. 6h-j), similarly to DMM, DES and LPS (Fig. 5a-e, 5p).

## Discussion

Glucocorticoid production in response to inflammation is essential for survival. The adrenal gland shows great resilience to damage induced by inflammation due to its strong regenerative capacity [7, 59,60]. This maintains glucocorticoid release during infection or sterile inflammation, which is vital to restrain and resolve inflammation [1,4]. However, severe sepsis is associated with adrenocortical impairment [11–15,61]. We show here that the inflamed adrenal cortex undergoes cellular metabolic reprograming which involves perturbations in the TCA cycle and oxidative phosphorylation, finally leading to impaired steroidogenesis. Our findings provide a mechanistic explanation of inflammation-related impaired adrenocortical steroidogenesis through cell metabolic reprogramming of steroidogenic adrenocortical cells.

In specific, we demonstrate that IL-1β reduces *SDHB* expression through DNMT1-dependent DNA methylation of the *SDHB* promoter. Several studies have shown that inflammation promotes DNA methylation and thereby regulates gene expression [62–66]. Particularly IL-1β was demonstrated to increase DNA methylation in different genes in a cell type-specific manner [63,67]. In accordance, DNMT1 expression was shown to increase upon acute inflammation in human peripheral blood mononuclear cells or mouse spleens [62,68], as well as in fibroblasts treated with IL-1β [67]. Moreover, reduced *SDH* promoter methylation associates with enhanced SDHB expression and reduced succinate levels in villi from individuals with recurrent spontaneous abortion [69]. These reports stand in accordance with our findings showing regulation of SDHB expression through its promoter methylation by an IL-1β – DNMT axis in steroidogenic adrenocortical cells. In contrast, itaconate, which was shown to reduce SDH activity in macrophages [54], does not regulate SDH in adrenocortical cells.

Consequently, accumulation of succinate leads to impaired oxidative phosphorylation and ATP synthesis, coupled to reduced steroidogenesis. Intact mitochondrial membrane potential and ATP generation are essential requirements for steroidogenic function [52,53]. This we confirmed by treatment of adrenocortical cells with the mitochondrial uncoupler FCCP or the ATP synthase inhibitor oligomycin, which both diminished steroidogenesis. Interestingly, a switch from the canonical towards a non-canonical TCA cycle, involving metabolism of mitochondrially derived citrate to acetyl-CoA, was recently described and may be activated in inflammation [70,71]. Whether a shift to the non-canonical TCA cycle regulates steroidogenesis remains to be elucidated.

SDH regulates ETC-mediated ROS formation: SDH inhibition or increased succinate levels augment ROS generation in tumors and macrophages [45,72–75]. Similarly, we show in adrenocortical cells that SDH inhibition or high succinate levels lead to increased ROS levels at the expense of mitochondrial oxidative function and ATP production. Adrenocortical disorders such as triple A syndrome and familial glucocorticoid deficiency can be driven by increased oxidative stress in the adrenal cortex [46]. In fact, mutations in genes encoding for proteins conferring antioxidant protection were implicated in the development of adrenocortical deficiencies [46]. Hence, SDH dysfunction leading to oxidative stress may be an important component of the pathophysiology of adrenocortical insufficiency, a notion which merits further investigation.

In conclusion, we demonstrate that tight regulation of succinate levels is essential for normal steroidogenesis, while disruption of SDH expression in inflammation through the IL-1β-DNMT1 axis contributes to adrenocortical dysfunction (Fig. 7). This study expands the current knowledge on regulation of glucocorticoid production and identifies potential targets for therapeutic interventions.

## Acknowledgements

We thank Christine Mund, Denise Kaden and Catleen Conrad for technical assistance. We acknowledge the technical support from the Core Facility Cellular Imaging of the Medical Faculty Carl Gustav Carus, TU Dresden for confocal imaging, and from the Light Microscopy Facility of the CMCB Technology Platform at TU Dresden, for laser microdissection.

## Statements & Declarations

### Author contributions

I.M.: planned and performed experiments, analyzed data, prepared the figures, wrote the manuscript; A.W: planned and performed experiments, analyzed data, edited the manuscript; E.H.: performed experiments, analyzed data; C.Y.: performed experiments; A.S.: analyzed data; E.T.: performed experiments; N.S.: performed experiments, analyzed data; O.K.: performed experiments; M.P.: performed experiments; H.A.: performed experiments; H.H.: edited the manuscript; W.K.: performed experiments; B.W.: edited and discussed the manuscript; C.T.: planned experiments; A.D.: planned experiments; A.W.: planned experiments; K.W.L: planned experiments; M.P.: analyzed data; T.C.: edited and discussed the manuscript; V.I.A.: conceived the study, supervised the project, wrote and edited the manuscript.

### Data Availability

The datasets generated during the current study are available from the corresponding author. RNA-Seq data are available in https://www.ncbi.nlm.nih.gov/geo/query/acc.cgi?acc=GSE200220

### Ethics approval

The animal experiments were approved by the Landesdirektion Sachsen Germany.

