## Supplemental Figures for "Succinate mediates inflammation-induced adrenocortical dysfunction"

***Correspondence**:

### Supplemental Figures

#### Suppl. Fig. 1

Efficiency of $CD31^{-}CD45^{-}$, immune ($CD45^{+}$) and endothelial ($CD31^{+}$) cell sorting. mRNA expression of *Star* (**a**), *Cd31* (**b**) and *Cd45* (**c**) in sorted $CD31^{-}CD45^{-}$, $CD45^{+}$ and $CD31^{+}$ cell populations from adrenal glands of mice 6 h post-injection of PBS or LPS (n = 6 mice per group). Data are presented as mean ± s.d., p values were calculated with Mann-Whitney U-test**.** **p < 0.01.

#### Suppl. Fig. 2

Inflammation-associated changes in the steroidogenic pathway. **a-e** mRNA expression of steroidogenic enzymes (*Star*, *Cyp11b1, 3β-Hsd2*, *Cyp21a1* and *Cyp11a1*) in adrenocortical cells of mice treated for 6 h with PBS or LPS (n = 6-8 mice per group, shown is one from two experiments). (**f**) Representative immunofluorescence images of adrenal gland sections from PBS and LPS mice stained for SF-1 (magenta) and DAPI (blue). Scale bar, 300 μm. Quantification of the mean fluorescence intensity of SF-1 staining in the adrenal cortex (excluding the outer capsule region) (n = 6 mice per group). Data are presented as mean ± s.d., p values were calculated with Mann-Whitney test**.** *p < 0.05, **p < 0.01.

#### Suppl. Fig. 3

Expression of TCA cycle genes in endothelial and immune cells of adrenal glands of LPS-treated mice. Expression of *Idh1*, *Idh2*, *Sdhb* and *Sdhc* in endothelial ($CD31^{+}$) (**a**) and immune ($CD45^{+}$) cells (**b**) sorted from adrenal glands of mice treated for 6 h with PBS or LPS (n = 6-8 mice per group, one from two experiments). Data are presented as mean ± s.d., p values were calculated with Mann-Whitney test**.** *p < 0.05, **p < 0.01, ***p < 0.001.

#### Suppl. Fig. 4

High succinate levels impair glucocorticoid production in adrenocortical cells. **a**,**b** NCI-H295R cells were treated for 24 h with DMM or DES and for another 24 h with Forskolin (Fsk) (n = 6). **c** NCI-H295R cells were transfected with si*SDHB* or control siRNA (siCtrl) and 24 h post-transfection they were treated for 24 h with Forskolin (n = 4). **d-f** NCI-H295R cells were treated for 24 h with the indicated concentrations of DES and for another 24 h with Forskolin (n = 4). Measurements for indicated steroid hormones were performed in supernatants of NCI-H295R cultures by LC-MS/MS. Data are shown as mean ± s.d., p values were calculated with one-way ANOVA (**a-c**) or two-way ANOVA (**d-f**). *p < 0.05, **p < 0.01, ***p < 0.001, ****p < 0.0001. BLD = below level of detection.

**Suppl. Fig. 5**

SiRNA silencing efficiencies. **a** Western blot analysis for SDHB in NCI-H295R cells transfected with 10, 30 or 50 nM si*SDHB* or siCtrl (24 h post-transfection). $\alpha$-TUBULIN was used as loading control. **b**,**c** mRNA expression of *Sdhb* and *Idh2* in primary adrenocortical cells transfected for 24 h with 10, 30 or 50 nM si*Sdhb*, si*Idh2* or siCtrl (n = 3). **d** *Dnmt1* expression in primary adrenocortical cells transfected for 24 h with 30 nM si*Dnmt1* or siCtrl (n = 5). Data are shown as mean ± s.d., p values were calculated with two-way ANOVA. $^*p < 0.05$, $^{**}p < 0.01$, $^{****}p < 0.0001$. Red boxes mark the concentrations with most efficient knock-down, which were chosen for further experiments.

**Suppl. Fig. 6**

Disruption of IDH function does not affect glucocorticoid production. **a**,**b** Primary adrenocortical cells were treated for 24 h with AG221 or DMSO and for another 45 min with ACTH (n = 6). **c,d** Adrenal gland explants were treated for 24 h with AG221 or DMSO and for another 45 min with ACTH (n = 4). **e**,**f** NCI-H295R cells were treated for 24 h with AG221 or DMSO and for another 24 h with Forskolin (n = 4-6). **g**,**h** Primary adrenocortical cells were transfected with si*Idh2* or siCtrl and 24 h post-transfection they were treated for 45 min with ACTH (n = 7). Measurements of steroid hormones were performed in supernatants of primary adrenocortical cell cultures, adrenal gland explants or NCI-H295R cells by LC-MS/MS. Data are shown as mean ± s.d. and analyzed with one-way ANOVA. $^*p < 0.05$, $^{**}p < 0.01$. BLD = below level of detection.

**Suppl. Fig. 7**

Itaconate does not affect SDH activity or steroidogenesis in adrenocortical cells. **a** mRNA expression of *Irg1* in $CD31^-/CD45^-$ and $CD45^+$ cells sorted from adrenal cortex of mice treated for 6 h with PBS or LPS (n = 8 mice per group). **b**,**c** Itaconate levels in $CD31^-/CD45^-$ (**b**) and $CD31^+/CD45^+$ cells (**c**) sorted from adrenal glands of mice treated for 24 h with PBS or LPS (n = 8 mice per group, shown is one from two experiments). **d-g** Itaconate, succinate and fumarate levels and succinate/fumarate ratio in lysates of primary adrenocortical cells treated for 24 h with 4-OI (n = 4). **h,i** Corticosterone and 11-deoxycorticosterone levels in the supernatant of primary adrenocortical cells treated for 24 h with 4-OI (n = 4). **j** Quantification of SDH activity in the adrenal cortex of *Irg1*-KO and wild-type mice treated for 16 h with LPS (n = 6 mice per group). Values are normalized to the total protein amount in the adrenal cortex. Data are shown as mean ± s.d. and analyzed with Mann-Whitney test. $^*p < 0.05$, $^{***}p<0.001$.

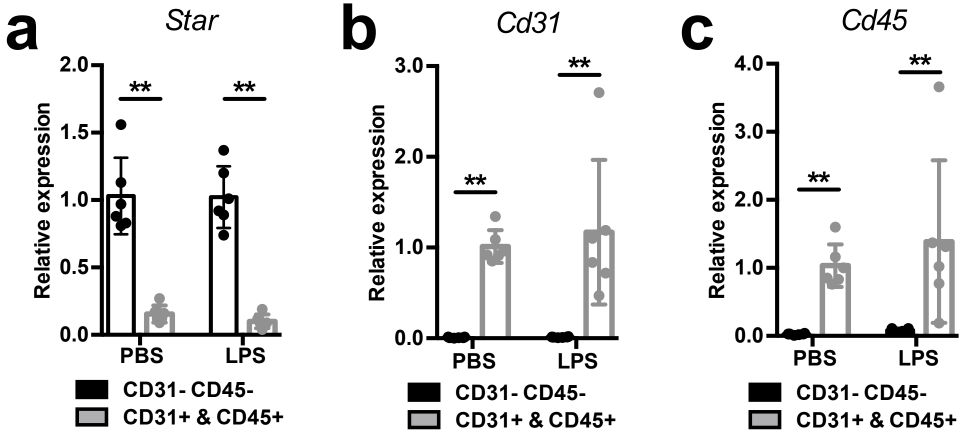

a
Star
2.0
1.5
1.0
0.5
0.0
Relative expression
**
**
PBS
LPS
CD31- CD45-
CD31+ & CD45+
b
Cd31
3.0
2.0
1.0
0.0
Relative expression
**
**
PBS
LPS
CD31- CD45-
CD31+ & CD45+
c
Cd45
4.0
3.0
2.0
1.0
0.0
Relative expression
**
**
PBS
LPS
CD31- CD45-
CD31+ & CD45+


Suppl. Fig. 1

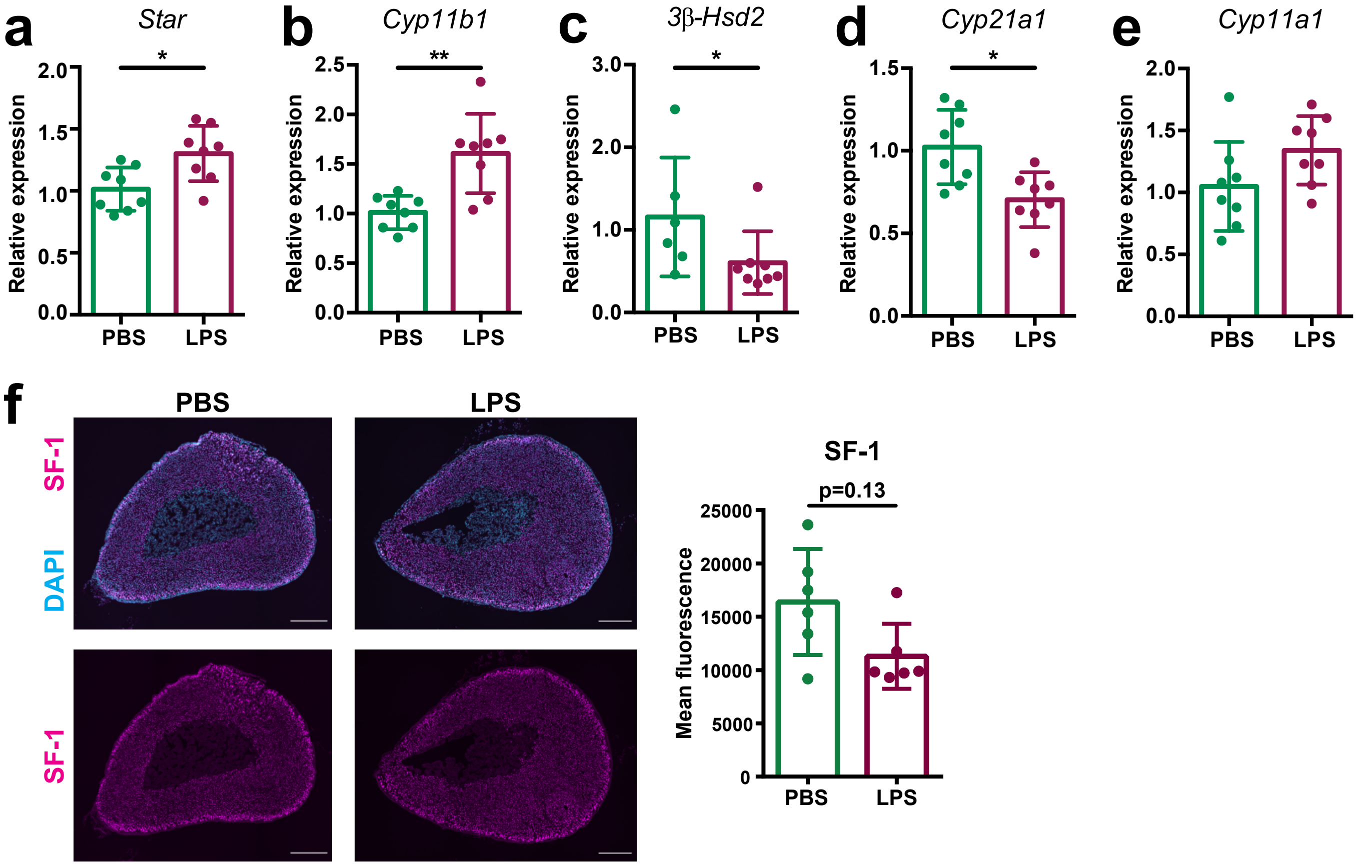
a
Star
Relative expression
2.0
1.5
1.0
0.5
0.0
*
PBS
LPS
b
Cyp11b1
Relative expression
2.5
2.0
1.5
1.0
0.5
0.0
**
PBS
LPS
c
3β-Hsd2
Relative expression
3.0
2.0
1.0
0.0
*
PBS
LPS
d
Cyp21a1
Relative expression
1.5
1.0
0.5
0.0
*
PBS
LPS
e
Cyp11a1
Relative expression
2.0
1.5
1.0
0.5
0.0
PBS
LPS
f
PBS
LPS
SF-1
DAPI
SF-1
SF-1
p=0.13
Mean fluorescence
25000
20000
15000
10000
5000
0
PBS
LPS

Suppl. Fig. 2

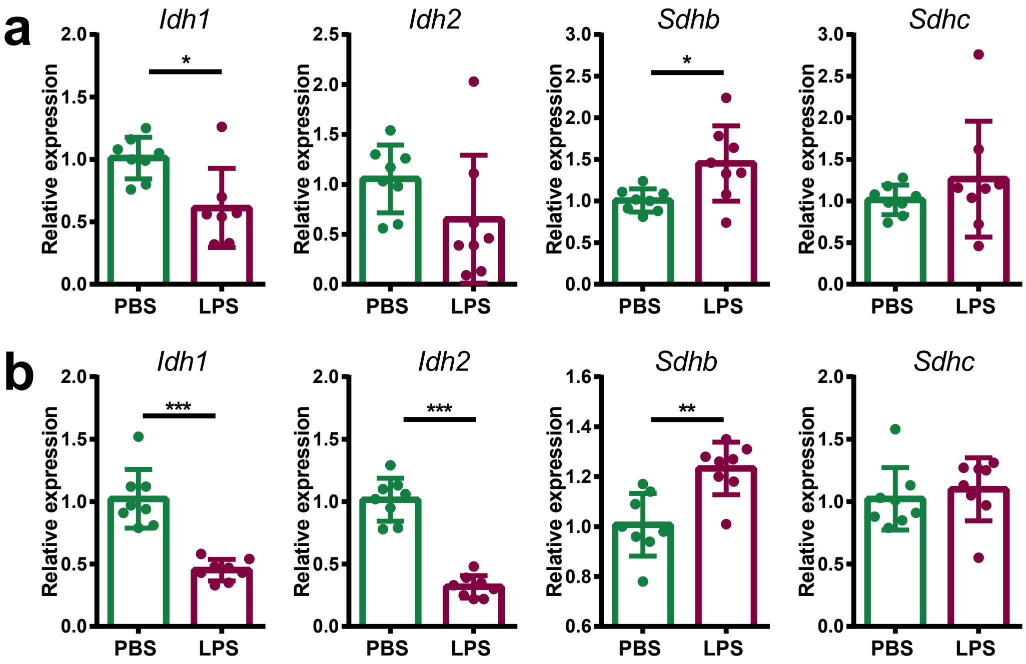


Suppl. Fig. 3

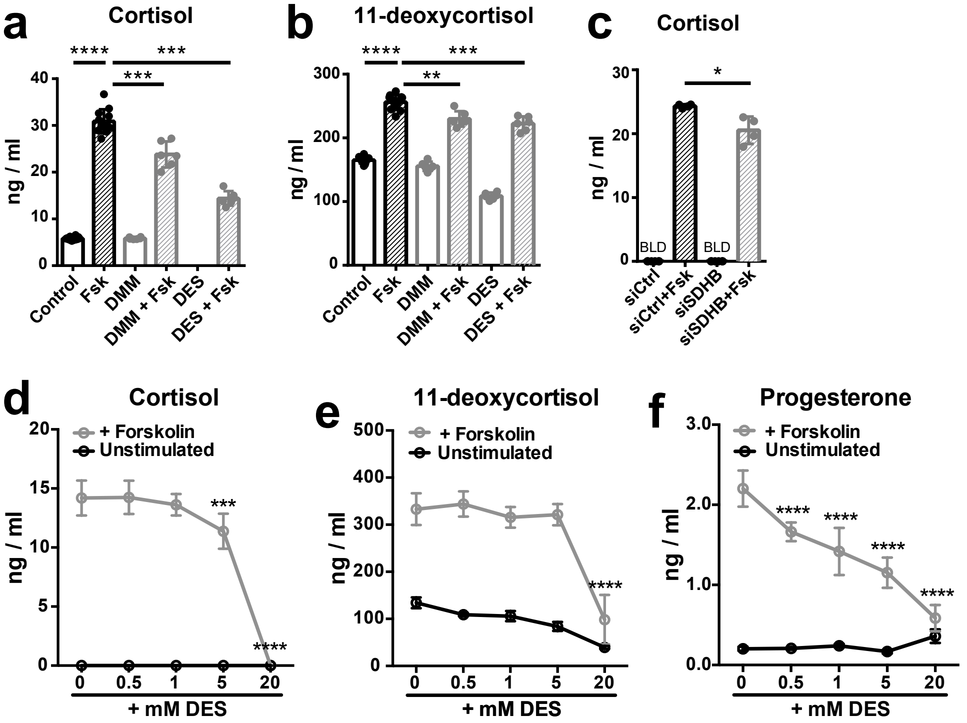
a
Cortisol
ng / ml
Control
Fsk
DMM
DMM + Fsk
DES
DES + Fsk
b
11-deoxycortisol
c
Cortisol
BLD
siCtrl
siCtrl+Fsk
siSDHB
siSDHB+Fsk
d
Cortisol
+ Forskolin
Unstimulated
+ mM DES
e
11-deoxycortisol
f
Progesterone

Suppl. Fig. 4

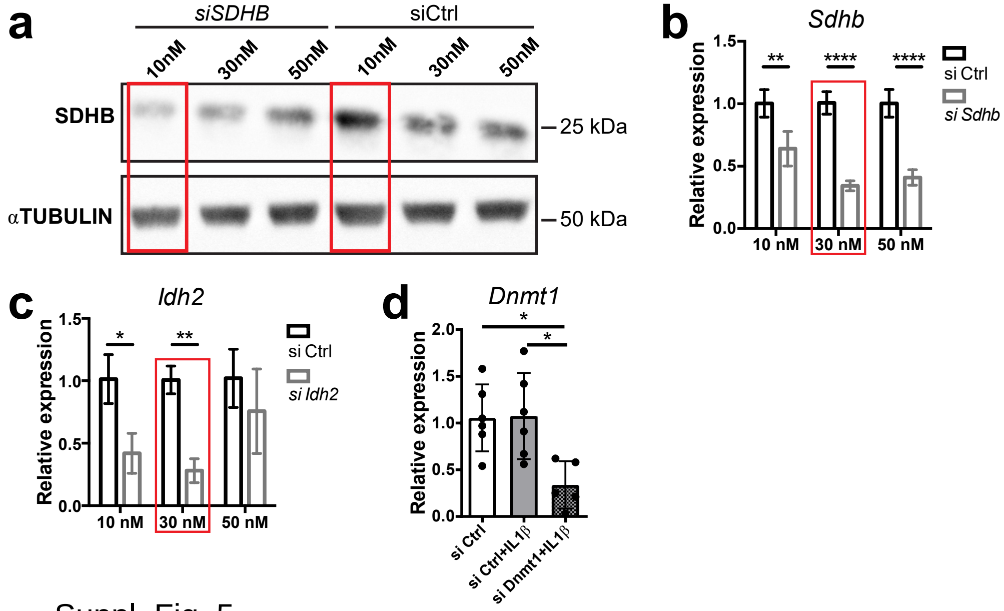
a
siSDHB
siCtrl
10nM
30nM
50nM
10nM
30nM
50nM
SDHB
αTUBULIN
25 kDa
50 kDa
b
Sdhb
Relative expression
1.5
1.0
0.5
0.0
**
****
****
10 nM
30 nM
50 nM
si Ctrl
si Sdhb
c
Idh2
Relative expression
1.5
1.0
0.5
0.0
*
**
10 nM
30 nM
50 nM
si Ctrl
si Idh2
d
Dnmt1
Relative expression
2.0
1.5
1.0
0.5
0.0
*
*
si Ctrl
si Ctrl+IL1β
si Dnmt1+IL1β

Suppl. Fig. 5

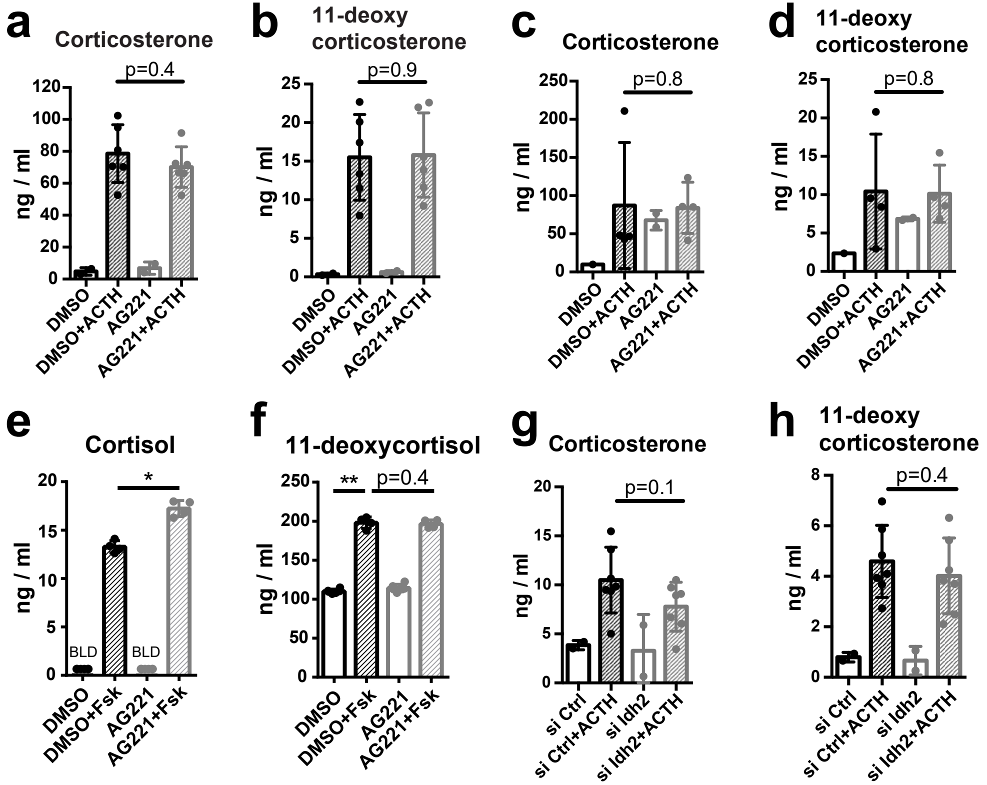
a
Corticosterone
p=0.4
ng / ml
DMSO
DMSO+ACTH
AG221
AG221+ACTH
b
11-deoxy corticosterone
p=0.9
c
Corticosterone
p=0.8
d
11-deoxy corticosterone
p=0.8
e
Cortisol
*
BLD
BLD
DMSO+Fsk
AG221+Fsk
f
11-deoxycortisol
**
p=0.4
g
Corticosterone
p=0.1
si Ctrl
si Ctrl+ACTH
si Idh2
si Idh2+ACTH
h
11-deoxy corticosterone
p=0.4

Suppl. Fig. 6

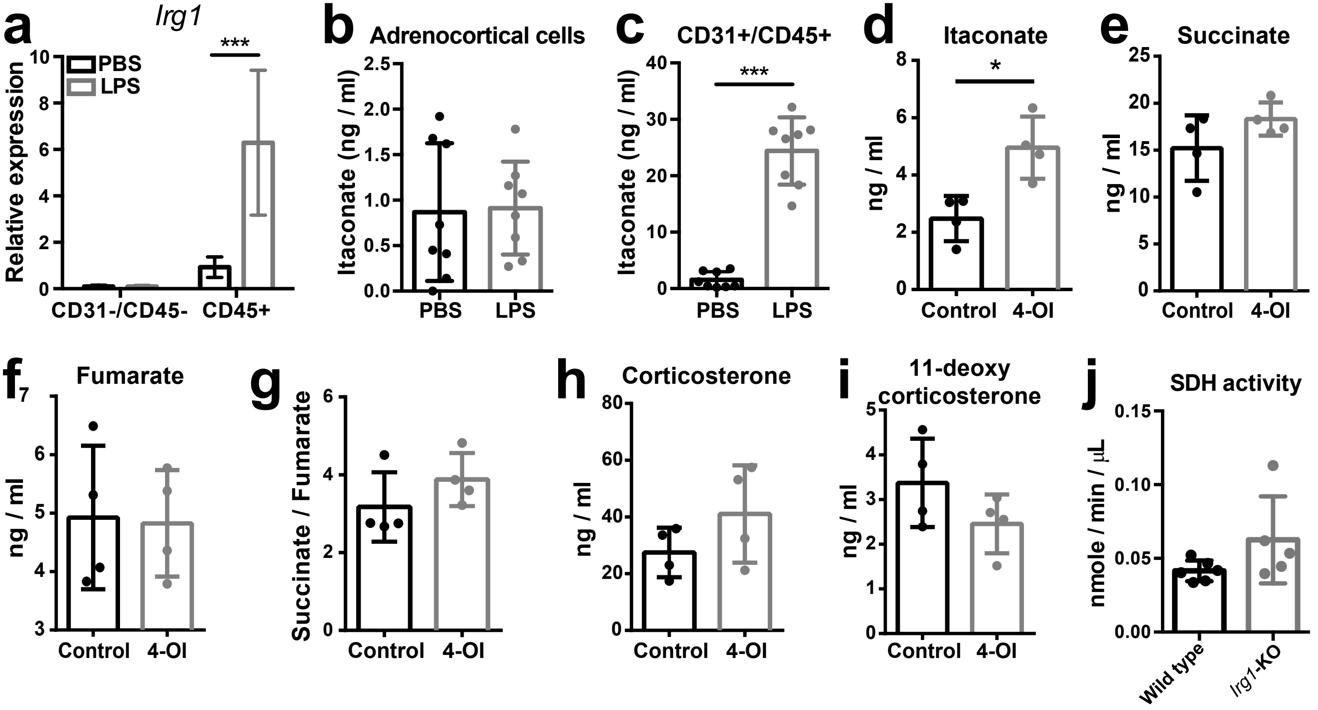
a
Irg1
***
PBS
LPS
Relative expression
CD31-/CD45-
CD45+
b
Adrenocortical cells
Itaconate (ng / ml)
PBS
LPS
c
CD31+/CD45+
***
Itaconate (ng / ml)
PBS
LPS
d
Itaconate
*
ng / ml
Control
4-OI
e
Succinate
ng / ml
Control
4-OI
f
Fumarate
ng / ml
Control
4-OI
g
Succinate / Fumarate
Control
4-OI
h
Corticosterone
ng / ml
Control
4-OI
i
11-deoxy corticosterone
ng / ml
Control
4-OI
j
SDH activity
nmole / min / μL
Wild type
Irg1-KO

Suppl. Fig. 7
